# Hidden molecular states of bacterial replicons beyond the chromosome–plasmid dichotomy

**DOI:** 10.64898/2026.08.31.748271

**Authors:** Shitong Zhong, Teng Wang

## Abstract

Bacterial genomes are organized into autonomous replicons, traditionally classified as either chromosomes or plasmids—a binary framework that underpins genome annotation and evolution models. Yet whether this binary framework captures the full diversity of replicon organization remains unclear. Here we show that bacterial replicons occupy three recurrent organizational states rather than two canonical categories. By integrating quantitative measures of chromosome–plasmid sequence affinity (plasmidness) across more than 72,000 replicons from 21 bacterial genera, we identify a distinct class—intermediate replicons—that occupies a positional and functional middle ground. These replicons are plasmid-sized, harbor substantial chromosomal sequence ancestry, and lack canonical replication signatures typically associated with either class. Multiple complementary molecular properties converge on this same state. Comparative genomic analyses reveal their enrichment near recurrent chromosome remodeling regions and reveal close evolutionary ties to conjugative and antimicrobial resistance plasmids. Metagenomic data further corroborate their presence across natural ecosystems. Together, these findings reveal a previously unrecognized replicon state and redefine bacterial genome organization beyond the chromosome–plasmid dichotomy.

## Introduction

Biological diversity rarely conforms to the boundaries we impose on it^1^. The traditional division of cellular life into prokaryotes and eukaryotes, for example, was reshaped by the discovery that the seemingly unified prokaryotic world encompasses fundamentally distinct molecular organizations represented by Bacteria and Archaea^1–3^. Such cases illustrate a recurring principle: when biological entities are defined by multiple underlying properties, binary classifications can conceal reproducible states that do not fit neatly into either category. The lesson is not that biological categories are unhelpful, but that their apparent completeness should not be assumed^4–6^.

Bacterial genomes are no exception. Autonomous replicons are conventionally divided into two canonical states: chromosomes, which provide the principal genomic backbone^7,8^, and plasmids, which replicate independently and often carry accessory functions^9–12^. This chromosome–plasmid dichotomy has been remarkably successful in genome annotation, comparative genomics and studies of horizontal gene transfer^11,13^. Its success, however, rests on a fundamental assumption: that every autonomous replicon can be assigned to one of these two organizational states^13–16^. Yet replicon identity is defined by multiple molecular properties—including sequence ancestry, replication architecture, functional organization and inheritance—that need not evolve in concert^13,17–19^. Whether these properties resolve into only two stable states, or reveal additional states obscured by binary classification, has remained unexplored.

Several observations already suggest that replicon identities are more fluid than conventional classification implies^13,14,20^. Plasmids can acquire chromosomal regions, while chromosomal segments can gain autonomous replication capacity and become independent genetic elements^21–24^. These examples demonstrate that chromosome and plasmid characteristics can become uncoupled during evolution^13,19,25^. What remains unclear is whether such uncoupling produces reproducible combinations of molecular features that define distinct replicon states, rather than representing isolated instances of genomic remodeling.

Addressing this question is challenging because existing replicon frameworks are designed primarily for binary classification rather than for discovering emergent organizational states^16,26^. Current approaches typically selected molecular features, including replication systems, nucleotide composition, conserved markers and sequence similarity, to assign replicons into predefined categories^13,26–28^. Although effective for typical chromosomes and plasmids, these categorical strategies provide limited resolution for replicons with discordant combinations of molecular properties. A quantitative framework integrating multiple dimensions of replicon organization is therefore needed to determine whether the chromosome–plasmid dichotomy captures the full landscape of autonomous replicon diversity or whether additional stable states remain hidden.

Here, by integrating sequence similarity, genomic architecture and multiple molecular features across more than 72,000 autonomous replicons from 21 bacterial genera, we establish a quantitative framework to resolve replicon organization beyond conventional chromosome– plasmid categories. We identify a previously unrecognized class of autonomous replicons that consistently occupies a distinct molecular state between canonical chromosomes and plasmids, which we term intermediate replicons (IRs). IRs are independently supported by multiple molecular dimensions, evade existing binary classification schemes, associate with recurrent chromosomal remodeling regions and occupy a close neighborhood to conjugative and antimicrobial resistance plasmids within plasmid sequence networks. Importantly, IRs are also detectable in metagenomic assemblies from diverse ecosystems, with particularly prominent representation in human-associated microbiomes, demonstrating that this organizational state extends beyond curated reference genomes. Together, these findings shift the view of bacterial replicons from discrete genomic categories to a quantitative landscape of molecular states, revealing hidden organizational principles that shape genome diversification and gene flow.

## Results

### A quantitative framework reveals a conserved intermediate state of bacterial replicons

To determine whether the chromosome-plasmid binary classification fully captures bacterial replicon diversity, we assembled a comprehensive dataset of 72,264 annotated replicons (30,061 chromosomes and 42,203 plasmids) from 28,581 complete bacterial genomes spanning 21 genera in the NCBI RefSeq database^29^. We then developed plasmidness, a quantitative measure of chromosomal versus plasmid sequence similarities that maps every replicon onto a common molecular coordinate system, independent of its existing annotation (Fig. 1A; Methods). Briefly, plasmidness quantifies, for each nucleotide position, the proportion of homologous sequences originating from plasmids relative to all genus-wide sequence matches, and averages these values across the entire replicon^30^. Consequently, plasmidness approaches zero for replicons whose sequences are predominantly chromosome-derived and approaches one for replicons sharing sequence similarity almost exclusively with plasmids.

**Figure 1.**
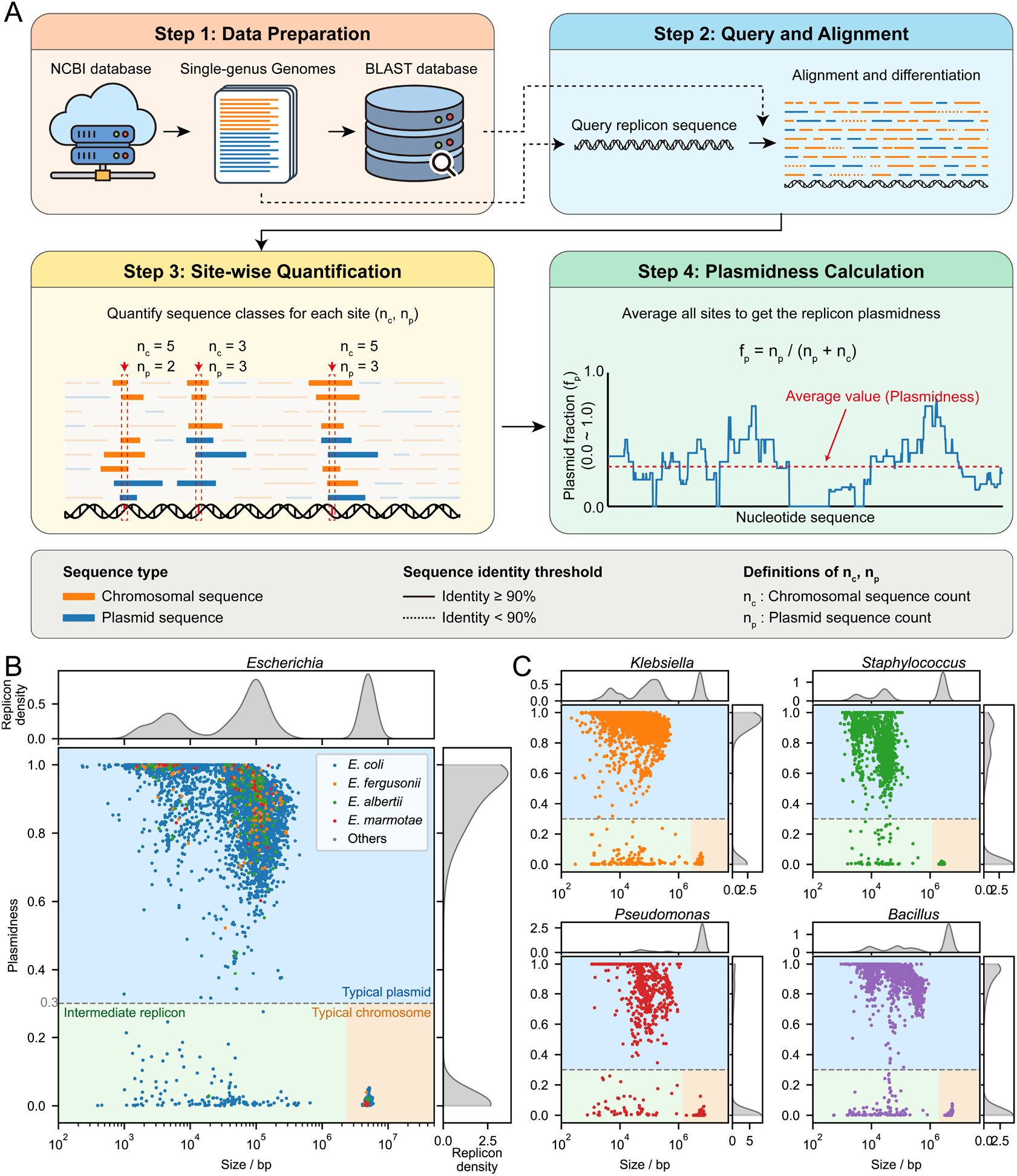
A quantitative replicon landscape reveals three conserved states across bacterial genera. (A) Schematic of the plasmidness framework. Genus-specific sequence similarity profiles were constructed across autonomous replicons, and the plasmidness score of each replicon was calculated from the genome-wide distribution of plasmid-associated sequence similarity. Higher plasmidness values indicate stronger sequence affinity to canonical plasmids. (B) Quantitative replicon landscape in *Escherichia*. Replicons occupy three discrete regions corresponding to typical plasmids, intermediate replicons (IRs), and chromosomes. Each point represents an individual replicon and is colored according to species origin. (C) Conservation of the three-state organization across bacterial genera. Replicon landscapes from four representative genera reveal the recurrent emergence of the same three molecular states across diverse bacterial lineages.

Applying this framework to *Escherichia* immediately revealed a previously unrecognized pattern of replicon organization. As anticipated, typical chromosomes and plasmids occupied two well-separated regions when plasmidness was plotted against replicon size (Fig. 1B). Chromosomes were uniformly large (>1 Mb) and exhibited near-zero plasmidness, whereas typical plasmids were substantially smaller and displayed high plasmidness, consistent with their limited sequence homology to host chromosomes.

Unexpectedly, however, a third, discrete population emerged. These replicons were comparable in size to typical plasmids yet exhibited plasmidness values indistinguishable from chromosomes, indicating that they retain extensive chromosomal sequence similarity despite much smaller sizes. Rather than forming a continuous gradient between chromosomes and plasmids, replicons clustered into three regions within this quantitative space, corresponding to typical chromosomes, typical plasmids, and a distinct intermediate state. To describe this previously unrecognized group without presupposing its evolutionary origin, we refer to them as intermediate replicons (IRs).

Remarkably, this three-state organization was not unique to *Escherichia*. Across all 21 bacterial genera examined, replicons reproducibly partitioned into the same three clusters despite substantial differences in genome architecture, plasmid abundance, and evolutionary history (Fig. 1C, Supplementary Figs. S1 and S2). Independent unsupervised hierarchical clustering^31^ of normalized replicon features further recovered the same three-state partition in most genera (Supplementary Fig. S3). The consistent emergence of IRs across phylogenetically diverse bacteria suggests that they represent a conserved mode of replicon organization rather than lineage-specific exceptions.

The three-state organization proved robust to analytical choices. Recomputing plasmidness using alternative BLAST similarity criteria, including analyses without alignment filtering or with a stringent 95% nucleotide identity threshold, produced nearly identical distributions and preserved the separation of the three replicon groups (Supplementary Fig. S4), demonstrating that IR identification is not dependent on specific parameter settings. In addition, the sequence alignments underlying plasmidness estimation were supported by extensive homology signals, with the number of matched sequences per site for IRs predominantly centered around an average magnitude of ∼10³ (Supplementary Fig. S5), indicating that IR assignment reflects broad sequence relationships rather than sparse or stochastic matches.

We next evaluated whether the identification of IRs could be explained by technical or assembly-related artifacts. IR-containing genomes showed no systematic bias across sampling year or sequencing approach, arguing against dataset-specific effects (Supplementary Fig. S6). Because IRs exhibit extensive chromosomal sequence similarity, we next examined whether they might represent misassembled chromosomal fragments. Highly similar IRs were independently detected across distinct genomes (Supplementary Fig. S7), and reassembly of representative genomes using alternative assembly strategies consistently recovered the same replicons with stable assembly graphs (Supplementary Fig. S8)^32–34^. Furthermore, several representative IRs have been reported in previous studies^35–38^, including pMTY18781-1_lncX3 (NZ_AP023206.1), which has been experimentally validated as an autonomous replicon^35^. Together, these evidences suggest that IRs represent authentic replicon entities rather than artifacts of genome assembly or sequencing processes.

### IRs challenge conventional chromosome–plasmid classification

The identification of IRs raises a fundamental question: does their current annotation as plasmids reflect a consistent biological identity or the limitations of existing classification criteria? Although most of IRs are annotated as plasmids in the NCBI RefSeq database^39^, their extensive chromosomal sequence ancestry suggest that they may not conform to the canonical properties used to define plasmids (Fig. 2A). We therefore examined whether other replicon classification frameworks provide consistent assignments for IRs.

**Figure 2.**
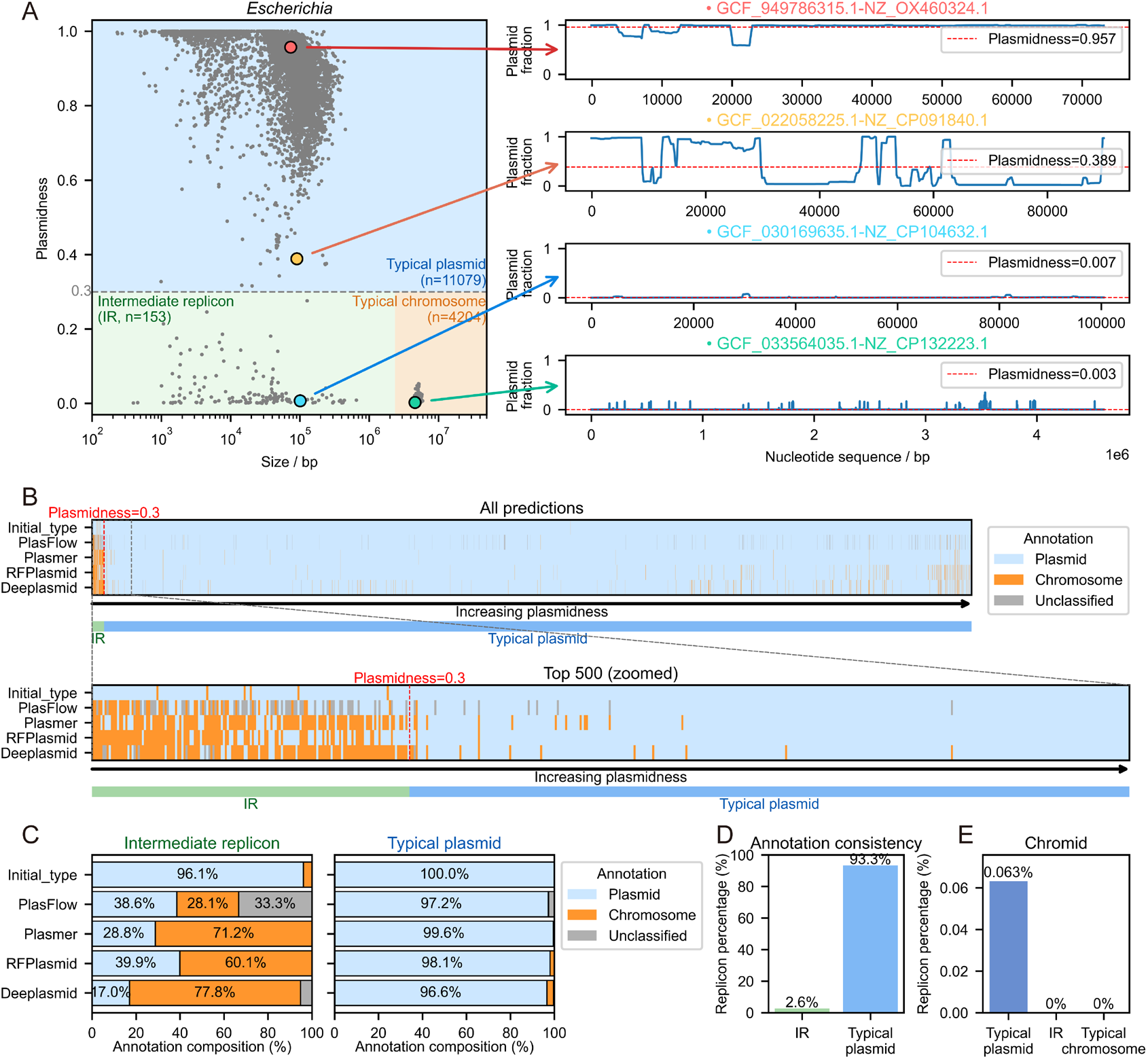
Intermediate replicons are not resolved by conventional replicon classification frameworks. (A) Plasmidness distributions across replicon classes. IRs occupy an intermediate region in plasmidness space despite displaying size distributions overlapping with typical plasmids, revealing discordance between conventional plasmid-associated features. (B) Classification of replicons using four independent prediction frameworks. Whereas typical plasmids show highly consistent assignments across methods, IRs exhibit substantial classification variability, with individual replicons frequently assigned to either plasmid or chromosome categories. (C) Predicted replicon assignments across classification methods. IRs display mixed classification outcomes, including a considerable fraction assigned as chromosomal by multiple approaches. (D) Cross-method agreement among replicon classifiers. IRs show significantly lower classification concordance than typical plasmids, highlighting their divergence from canonical replicon signatures. (E) Comparison with chromid annotations. Application of a chromid prediction framework identifies no overlap between IRs and annotated chromids in *Escherichia*, indicating that IRs represent a distinct replicon state.

To address this, we compared RefSeq annotations with four independent replicon classifiers based on distinct biological features, including nucleotide composition, replication systems and conserved marker genes^27,28,40,41^. As expected, canonical plasmids showed near-complete agreement across all methods, reflecting the robustness of existing classification schemes for typical replicons. In contrast, IRs displayed pronounced classification discordance, with identical replicons assigned to either plasmid or chromosome categories depending on the method used (Fig. 2B–D).

This classification discordance was observed across all bacterial genera examined (Supplementary Fig. S9). Because the evaluated approaches rely on distinct molecular criteria (PlasFlow^27^ relies solely on *k*-mer profiles; Plasmer^40^ and RFPlasmid^28^ further include marker genes, whereas Deeplasmid^41^ adopts deep neural networks to automatically extract sequence features without predefined genomic markers), their recurrent disagreement indicates that IRs deviate from the canonical feature combinations underlying chromosome or plasmid classification. Rather than reflecting methodological uncertainty, this ambiguity stems from the intrinsic architecture of IRs, which resists binary classification.

The distinct positioning of IRs relative to plasmids and chromosomes further prompted comparison with chromids, a recognized class of replicons characterized by chromosomal gene content and plasmid-derived maintenance and replication systems^14^. Applying chromid-finder^42^ to *Escherichia* replicons revealed no overlap between IRs and chromids (Fig. 2E). Most previously identified chromids occurred in *Vibrio* and *Burkholderia* and localized within the chromosome-like region of our quantitative framework, whereas IRs formed a distinct cluster (Supplementary Fig. S10).

Together, these findings demonstrate that IRs are not adequately represented by existing chromosome–plasmid classifications or by the current chromid framework. More broadly, they reveal that categorical annotation can obscure replicons with distinct molecular architectures when complex genomic variation is forced into predefined classes. Quantitative mapping of replicon states therefore provides a complementary view of bacterial genome organization beyond conventional classification.

### Multiple molecular dimensions define the intermediate replicon state

The unique positioning of IRs in replicon sequence space raises the question of whether this organization reflects a biologically meaningful genomic configuration. We therefore characterized IRs across multiple dimensions, including replication architecture, functional organization, sequence composition, evolutionary constraint, and replication-associated features, to determine whether their molecular properties converge on a coherent replicon state.

We first examined replication-associated features. As expected, chromosomes frequently contained *oriC*-associated sequences^43,44^, whereas typical plasmids carried plasmid-specific *oriV* systems^45^ (Fig. 3A, Supplementary Fig. S11). In contrast, approximately one-quarter of IRs contained putative *oriC*-associated regions but lacked detectable *oriV* elements (Fig. 3A). Consistently, while chromosomes uniformly encoded DnaA-like replication initiators and plasmids encoded plasmid-specific replication proteins^46,47^, IRs generally lacked recognizable initiators characteristic of either category (Fig. 3B; Supplementary Fig. S11). IRs were also depleted for canonical partitioning and conjugation systems^48–50^. Collectively, these findings point to a replication–maintenance architecture distinct from that of typical chromosomes or plasmids.

**Figure 3.**
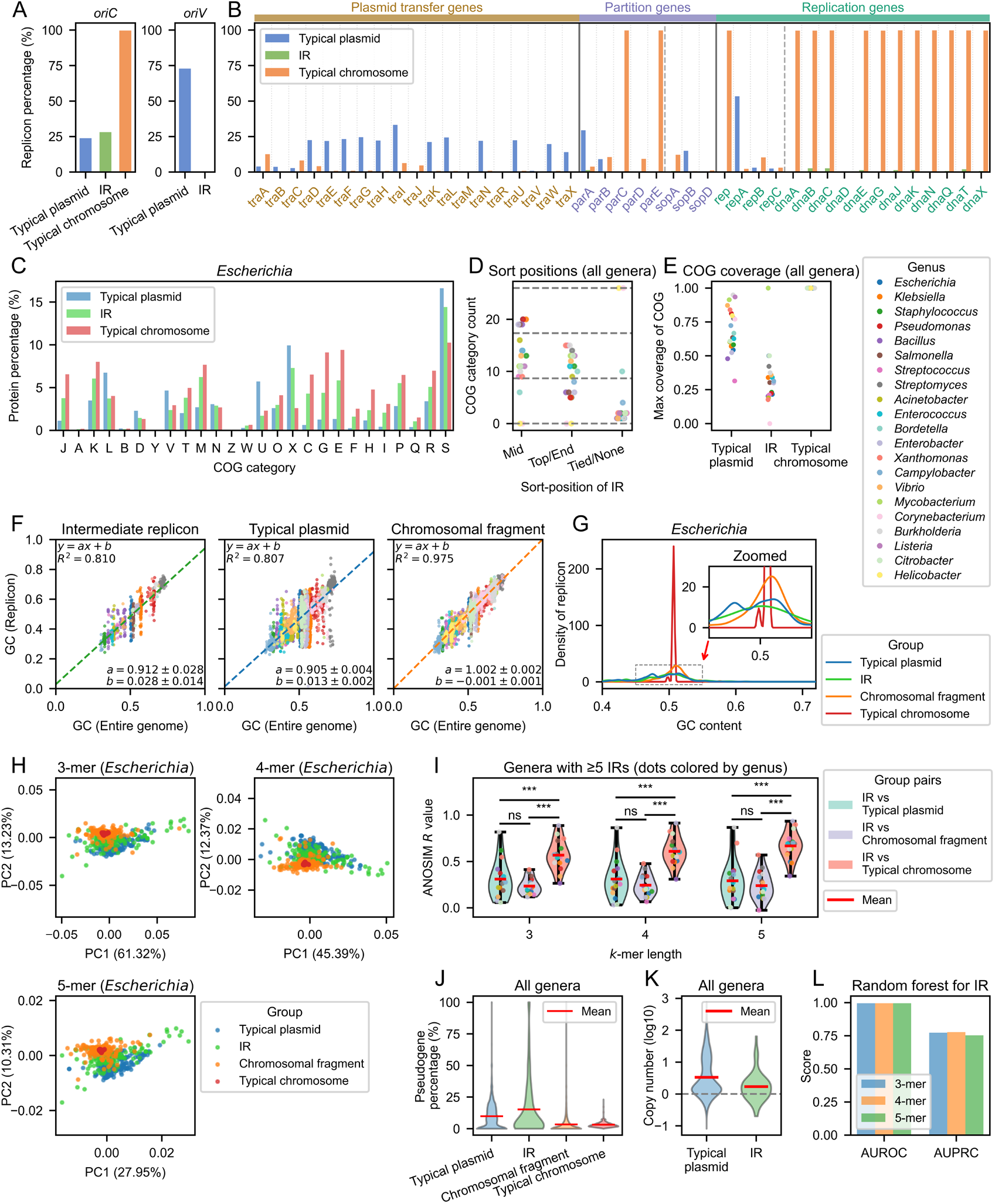
Intermediate replicons represent a coherent molecular state defined by multiple genomic dimensions. (A) Distribution of *oriC* and *oriV* sites across replicon classes. IRs are depleted in *oriV* sites and show reduced prevalence of *oriC* relative to chromosomes. (B) Prevalence of functional core-gene markers among replicon classes. IRs exhibit reduced prevalence of core-gene markers relative to typical plasmids and chromosomes. (C) Functional organization of replicons based on COG category composition in *Escherichia*. The y-axis (protein percentage) represents the proportion of protein counts for each category relative to the total number of proteins within the corresponding replicon group. IRs display a functional profile intermediate between chromosomes and typical plasmids, reflecting coordinated contributions from both replicon classes. (D) Conservation of intermediate functional organization across bacterial genera. COG-based functional profiles consistently position IRs between chromosomes and plasmids across diverse lineages. The x-axis denotes sort position, defined as the rank position of IRs when three replicon groups (typical plasmid, IR, typical chromosome) were sorted in descending order by protein percentage. Sort-position annotations: Top (IR ranked first), Mid (IR ranked second), End (IR ranked last), Tied (IR shared equal protein-percentage values with at least one other replicon group), and None (no IR data available for a given genus). Each genus was further quantified by counting distinct COG categories assigned to each sort-position class. (E) Distribution of conserved core COG repertoires. IRs lack the extensive conserved core gene sets characteristic of chromosomes, while remaining distinct from typical plasmids. For a given genus and replicon group, COG coverage for each COG identifier was calculated as the fraction of replicons harboring that COG within this replicon group; the maximum value among all COG identifiers was defined as max coverage of COG for that replicon group. (F) GC content distributions of replicons and chromosomal fragments. IRs exhibit GC profiles closer to plasmids than to chromosomal fragments. Uncertainties for the slope and intercept represent the half-widths of the 95% confidence intervals. (G) GC content distributions in *Escherichia*. IRs show distribution patterns close to those of plasmids. (H) Sequence composition landscape based on *k*-mer profiles in *Escherichia*. Principal-component analysis reveals distinct sequence organization among chromosomes, chromosomal fragments, typical plasmids and IRs, with IRs occupying an intermediate region between canonical replicon classes. PC1 and PC2 denote the first and second principal components; the percentage values on each axis represent the proportion of total variance explained by that principal component. (I) ANOSIM-based similarity comparisons across replicon classes. IRs show highest similarity to chromosomal fragments. Pairwise comparisons between replicon classes were additionally performed using Welch’s t-test (two-sided). Significance markers: ns, not significant; *, *p* < 0.05; **, *p* < 0.01; ***, *p* < 0.001. (J) Pseudogene content across replicon types. IRs exhibit elevated pseudogene percentage relative to plasmids and chromosomes. Pseudogene percentage was calculated for each individual replicon or selected sequence as the fraction of pseudogene CDS among all CDS features from GBFF records. (K) Copy number distribution across replicons. IRs display intermediate copy numbers between plasmids and chromosomes, with values slightly above unity. (L) Computational identification of IRs using integrated molecular features. A random-forest classifier trained on COG composition and *k*-mer features distinguishes IRs from canonical replicon classes, demonstrating that their molecular identity is encoded across multiple genomic dimensions. AUROC, area under the receiver operating characteristic curve; AUPRC, area under the precision– recall curve.

We next asked whether the unique identity of IRs is reflected in their functional organization. COG-based profiling revealed clear differences between chromosomes and typical plasmids, with chromosomes enriched for conserved core functions and plasmids dominated by accessory functions (Fig. 3C; Supplementary Fig. S12)^51^. In contrast, IRs consistently displayed functional compositions intermediate between these two canonical replicon states across most COG categories, a pattern reproduced across all bacterial genera examined (Fig. 3C, D; Supplementary Fig. S12). Because gene-content composition can be influenced by replicon size, we performed principal-component analysis^52^ using randomly sampled chromosomal fragments matched to the size distributions of plasmids and IRs. IRs occupied a distinct intermediate position between chromosomal fragments and typical plasmids (Supplementary Fig. S13). At the level of individual COG families, IRs lacked a conserved core repertoire or defining functional markers (Fig. 3E)^51^. These findings indicate that IRs are characterized by an intermediate functional profile rather than by a specific set of marker genes.

We then examined whether IRs exhibit signatures of an independent evolutionary history. To control for the potential influence of replicon size, we generated random chromosomal fragments matched to the size distributions of plasmids and IRs. These fragments closely recapitulated the GC content of their source genome (R² = 0.975), indicating strong host-specific compositional coupling independent of sequence length. In contrast, typical plasmids and IRs showed substantially weaker correlations with host genome GC content (R² = 0.807 and 0.810, respectively) (Fig. 3F)^53^. Consistently, in *Escherichia*, IRs displayed GC distributions distinct from host chromosomes and more similar to typical plasmids (Fig. 3G). Thus, despite extensive chromosomal sequence ancestry, IRs are partially decoupled from host chromosome composition, suggesting that they undergo independent compositional evolution rather than representing chromosomal fragments.

This distinction extended beyond GC content. Analysis of oligonucleotide frequencies (*k* = 3–5) revealed separate sequence signatures for chromosomes and typical plasmids^54,55^, whereas IRs occupied a reproducible region between these two states (Fig. 3H). Consistently, calculations of Analysis of Similarities (ANOSIM) *R* values^56^ across bacterial genera showed that IRs exhibited comparable compositional distances from chromosomal fragments and plasmids, rather than preferential similarity to either class (Fig. 3I). These results indicate that IRs possess a distinct sequence architecture incorporating features of both canonical replicon types.

We next investigated whether IRs experience distinct evolutionary constraints. Chromosomes and size-matched chromosomal fragments showed uniformly low pseudogene frequencies^57^, consistent with strong purifying selection on conserved genomic functions, whereas typical plasmids accumulated higher pseudogene densities (see Methods for more details). Notably, IRs exhibited the highest pseudogene frequencies among all replicon states (Fig. 3J, Supplementary Fig. S14A), indicating elevated gene turnover and relaxed selective constraint relative to both canonical chromosomes and plasmids.

Finally, we examined whether this distinct evolutionary organization was reflected in replication dynamics. Using experimentally derived plasmid-to-chromosome copy-number ratios^58^, we found that typical plasmids displayed broad distribution in copy number, whereas IRs consistently maintained low but non-unit copy-number ratios, generally ranging from one to ten copies per chromosome (Fig. 3K; Supplementary Fig. S14B). These patterns indicate that IRs undergo limited replication amplification, distinct from both chromosomal maintenance and high-copy plasmid replication.

Comparison with chromids revealed clear molecular separation between the two groups (Supplementary Fig. S15)^14,42^, indicating that IRs do not correspond to a previously recognized replicon category. Together, these independent dimensions converge on a coherent picture: IRs represent a reproducible replicon state characterized by coordinated differences in sequence evolution, selective constraint and replication strategy. Their identity cannot be explained by any single molecular feature or by partial resemblance to either chromosomes or plasmids. Instead, replicon states emerge from the integration of multiple evolutionary and molecular constraints.

We next asked whether the distinct molecular characteristics of IRs are sufficient to computationally distinguish them from canonical replicon classes. We first examined whether genomic foundation model embeddings could separate IRs from chromosomes and typical plasmids without predefined labels. Across several genomic foundation models (Evo2^59^, gLM2^60^ and Nucleotide Transformer^61^), unsupervised embedding projections showed limited separation of IRs from canonical replicon classes (Supplementary Fig. S16), indicating that general sequence representations alone do not fully capture the multidimensional features defining IR identity. We therefore trained supervised classifiers using the integrated molecular features (COG category frequencies and *k*-mer profiles). A random-forest model achieved robust discrimination of IRs from chromosomes and typical plasmids (Fig. 3L; Supplementary Fig. S17), demonstrating that IRs possess a reproducible computationally identifiable signature arising from multiple genomic dimensions.

### Intermediate replicons reveal structured chromosome–replicon remodeling regions

The extensive chromosome-like sequence ancestry of IRs raises the question of whether their chromosome associations occur randomly across bacterial genomes or preferentially involve specific genomic regions. If such interactions were random, homologous sequences should be broadly scattered across chromosomes. Alternatively, recurrent chromosome– replicon exchange would be expected to generate preferential associations with discrete genomic loci.

To distinguish between these possibilities, we mapped high-confidence sequence homology between replicons—including IRs and typical plasmids—and chromosomes across *Escherichia* genomes (Fig. 4A). Unlike typical plasmids, which showed limited chromosomal associations, IRs exhibited extensive homology with multiple chromosomal regions across diverse lineages. Among analyzed genomes, one chromosome (GCF_022569795.1)^29^, which contained the largest number of homologous associations, provided sufficient resolution to examine the spatial organization of replicon-associated regions. This chromosome exhibited homologous segments with 48 IRs and 1,424 typical plasmids.

**Figure 4.**
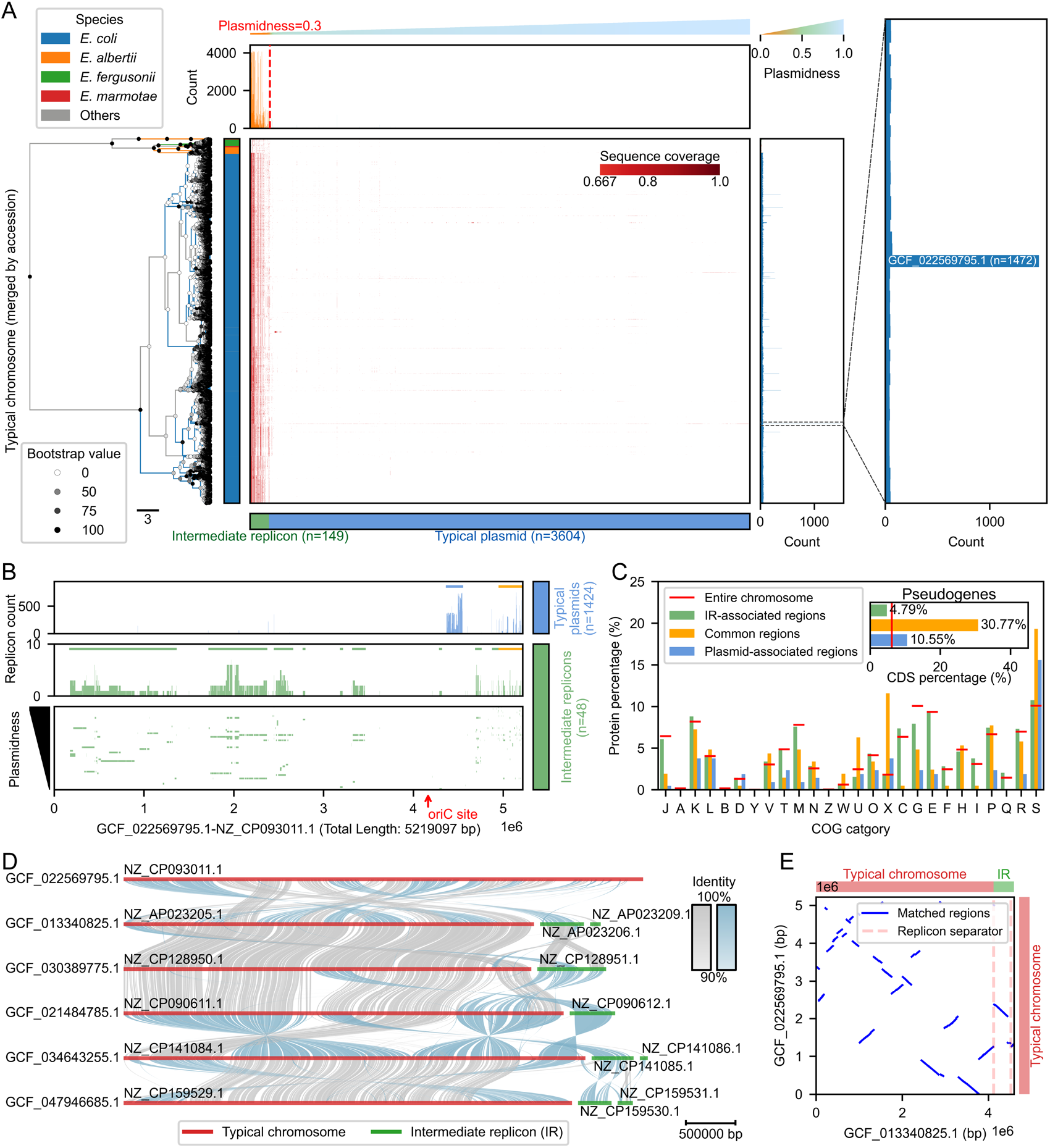
Intermediate replicons reveal structured chromosome–replicon associations in *Escherichia*. (A) Genome-wide association matrix between chromosomes and other replicon types based on shared homologous sequences. Chromosomes show significantly more associations with IRs than with typical plasmids. A reference chromosome (genome accession: GCF_022569795.1) exhibits the highest connectivity to both replicon classes. (B) Spatial distribution of replicon-associated regions along the reference chromosome. IR-associated sequences map to multiple distributed chromosomal loci, whereas typical plasmid-associated regions are concentrated within a smaller number of genomic regions. A shared hotspot (yellow) is associated with both replicon classes. (C) Molecular characteristics of chromosome-associated regions. Regions associated with IRs retain functional compositions similar to the chromosomal background, whereas plasmid-associated regions display distinct functional repertoires and elevated pseudogene content. Protein percentage refers to the proportion of proteins with valid amino-acid translation sequences within each genomic region. Pseudogene percentage was calculated per genomic region as the fraction of pseudogene CDS among all CDS features within that region from GBFF records. (D) Representative examples of homologous sequence associations between IRs and chromosomal loci across independent genomes. (E) Comparative genome alignment illustrating structural variation associated with IR-related regions. Genome comparisons reveal local rearrangements surrounding IR-associated loci, consistent with recurrent chromosome–replicon remodeling.

Projection of homologous regions onto this reference chromosome revealed different association patterns between replicon states (Fig. 4B). Typical plasmids were predominantly associated with a limited number of localized chromosomal regions, whereas IRs corresponded to more dispersed loci distributed across the chromosome. Despite these differences, both replicon classes converged on a shared hotspot region (Fig. 4B), indicating that specific chromosomal loci are repeatedly associated with autonomous replicons.

We next examined the molecular properties of these recurrently associated regions. Chromosomal loci preferentially linked to IRs retained functional compositions closely resembling the chromosomal background, whereas plasmid-associated regions exhibited markedly distinct functional repertoires (Fig. 4C, Supplementary Fig. S18). Notably, the shared hotspot was enriched for mobilome-related functions (COG category X), including prophage- and transposon-associated genes, and exhibited the highest pseudogene density among all analyzed regions. These features identify mobilome-rich genomic regions as candidate hotspots of chromosome–replicon remodeling.

Comparative analyses of representative genomes further revealed that IR-associated regions frequently coincide with local chromosomal rearrangements (Fig. 4D, E)^62^. Different IRs corresponded to distinct chromosomal segments, and their associated genomic contexts showed extensive structural variation across genomes. These observations suggest that IR-associated loci are embedded within dynamic genomic regions undergoing recurrent remodeling.

To identify genomic features associated with these remodeling-prone regions, we compared genes surrounding IR–chromosome homologous boundaries with randomly selected genomic regions. Genes associated with COG category X were significantly enriched near IR-associated boundaries (Supplementary Fig. S19), most of which corresponded to phage- and transposon-related functions^63^. Thus, mobile genetic element-associated regions appear preferentially associated with IR remodeling boundaries.

Collectively, these analyses reveal that chromosome–replicon associations are structured rather than randomly distributed. IRs preferentially associate with specific genomic regions enriched in mobile genetic elements and structural variation, suggesting that mobilome-rich loci represent recurrent environments for replicon remodeling. These observations define a genomic landscape associated with IR dynamics without implying a single direction or mechanism of replicon transition.

### IRs occupy a distinct network position linked to conjugative and antibiotic-resistant plasmids

Having established IRs as a distinct molecular state with extensive chromosomal ancestry, we next asked how they relate to the broader plasmid population. Specifically, we investigated whether IRs represent isolated replicon configurations or are preferentially connected to specific classes of plasmids through shared sequence relationships. To address this question, we constructed genus-specific sequence similarity networks in which individual plasmids or IRs were represented as nodes and edges connected replicons sharing extensive nucleotide homology (aligned regions spanning at least two-thirds of one sequence) (Fig. 5A; Methods). The *Escherichia* network comprised 11,059 plasmids or IRs linked by 1,742,174 similarity edges, enabling the global organization of replicon sequence space to be resolved.

**Figure 5.**
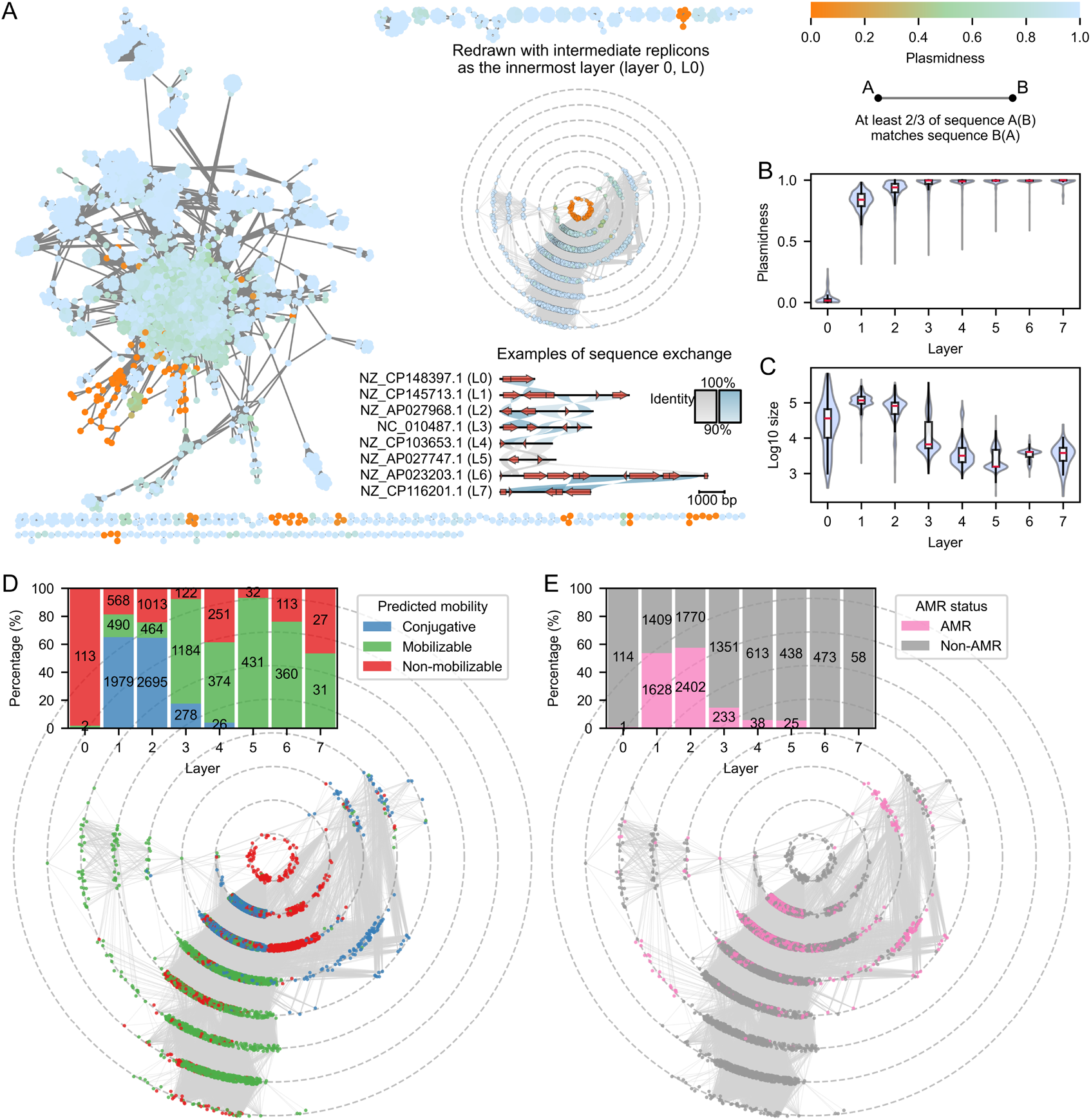
Intermediate replicons occupy a structured neighborhood within plasmid sequence space. (A) Sequence similarity network of IRs and typical plasmids. An edge was defined when the aligned regions covered at least two-thirds of either replicon. Replicons are reorganized into hierarchical network layers according to their shortest sequence-similarity distance from IRs (layer 0). IRs occupy a defined region within plasmid sequence space and are connected to surrounding plasmid populations through graded sequence similarity relationships. (B) Plasmidness variation across network layers. Replicons progressively increase in plasmid-like sequence characteristics with increasing topological distance from IRs. (C) Replicon size distribution across network layers. Network layers exhibit systematic shifts in replicon size, revealing structured variation across the sequence landscape. (D) Mobility profiles across network layers. Conjugative plasmids are preferentially enriched in the immediate sequence neighborhood of IRs and decline with increasing network distance. (E) Distribution of antimicrobial resistance-associated plasmids across network layers. AMR-associated plasmids show a similar enrichment pattern near IRs, identifying the local plasmid neighborhood surrounding IRs as a region of elevated mobility and resistance potential.

IRs were not randomly distributed in the network but instead occupied a highly structured neighborhood. We stratified typical plasmids according to their shortest network distance from IRs and examined how plasmid properties varied across successive network layers (Fig. 5B, C). A pronounced mobility gradient emerged around IRs. Plasmids directly connected to IRs (Layer 1) were strongly enriched for conjugative plasmids^64^, and this enrichment remained evident in Layer 2 before progressively declining at increasing network distances (Fig. 5D; Supplementary Fig. S20). A similar pattern was observed for antimicrobial resistance (AMR). Plasmids carrying AMR genes were likewise concentrated within the immediate neighborhood surrounding IRs and became progressively less frequent in more distant layers (Fig. 5E; Supplementary Fig. S20). These observations were robust across different coverage thresholds, with no substantial shifts in the overall patterns (Supplementary Fig. S21).

Notably, these sequence relationships do not imply evolutionary directionality. Similarity networks cannot distinguish whether IRs arise from mobile plasmids, contribute sequences to them, or undergo repeated reciprocal exchange. Instead, they reveal that IRs occupy a reproducible interface between chromosome-associated replicons and plasmid populations enriched for mobility and adaptive functions.

### Intermediate replicons occur across diverse natural microbial ecosystems

To determine whether IRs represent naturally occurring genomic entities beyond isolate-derived genomes, we examined their distribution in metagenomic datasets using the IMG/PR plasmid database^65^. IRs were detected across diverse ecological environments, demonstrating that this replicon state is not restricted to cultured bacterial genomes. Unlike isolate genomes, in which *Escherichia* contained the largest number of identified IRs, metagenome-derived IRs were particularly enriched in *Streptococcus* (Fig. 6A; Supplementary Fig. S22) and were predominantly recovered from human-associated environments (Fig. 6B). Genomic characterization further showed that most metagenomic IRs lacked detectable mobile genetic element signatures and antimicrobial resistance genes (Fig. 6B), consistent with the genomic properties observed in isolate-derived IRs.

**Figure 6.**
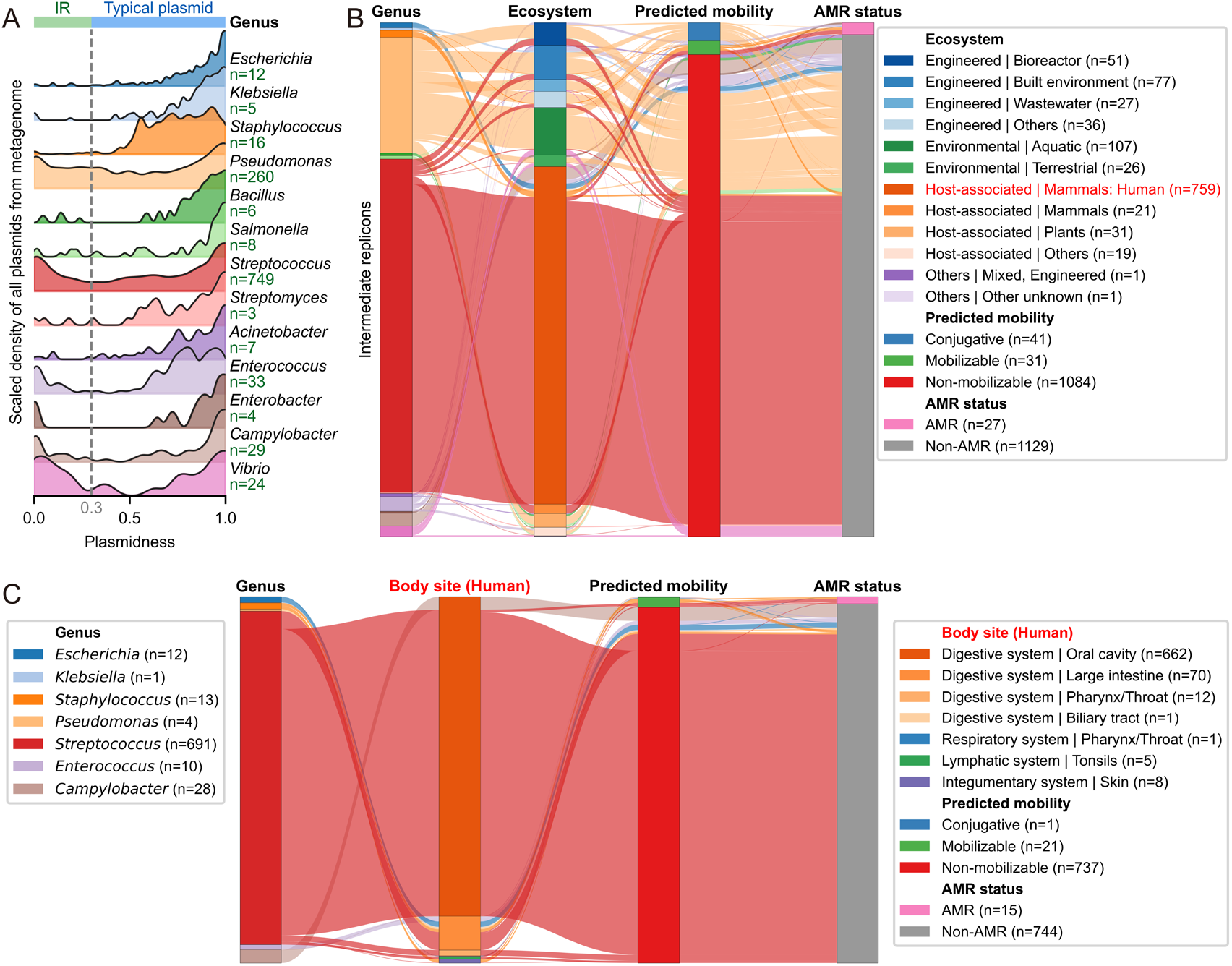
Intermediate replicons occur in natural microbial communities across diverse ecosystems. (A) Distribution of metagenome-derived replicons across replicon state space. Intermediate replicons are detected among metagenome-assembled plasmids from diverse bacterial genera. Density values were scaled 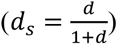 for visualization of low-abundance replicon states; only genera containing detectable IRs are shown. (B) Ecological distribution and functional characteristics of metagenomic IRs. IRs are enriched in human-associated environments, with a substantial proportion recovered from *Streptococcus*. Most IRs lack identifiable mobility-associated elements and antimicrobial resistance genes, consistent with their genomic characteristics in reference genomes. (C) Distribution of human-associated IRs across host-associated niches. Human-derived IRs show pronounced enrichment in the oral microbiome compared with other sampled body sites.

Within human-associated microbiomes, IRs showed pronounced ecological structuring, with the majority recovered from oral microbiomes rather than other sampled body sites, including the gut and skin (Fig. 6C). This distribution pattern suggests that IR occurrence is not random across host-associated environments and may reflect differences in microbial community composition, ecological interactions, or local selective pressures among distinct microbial habitats. Together, these observations demonstrate that IRs represent naturally occurring replicon states with non-uniform ecological distributions across microbial ecosystems.

## Discussion

For decades, autonomous bacterial replicons have been interpreted through a binary chromosome-plasmid framework. Our analyses suggest that this long-standing view might be incomplete. By quantifying multiple molecular dimensions, we identify a conserved IR state that is systematically overlooked by existing classification schemes yet reproducibly detected across diverse bacterial lineages. Rather than representing isolated anomalies or ambiguous annotations, IRs reveal a previously unrecognized level of organization within bacterial genomes. Notably, we screened complete plasmids derived from metagenomic samples within the IMG/PR plasmid dataset^65^ and successfully recovered large amounts of IRs (Fig. 6; Supplementary Fig. S22), demonstrating that this category of replicons is not rare in natural environmental niches.

The identification of IRs changes how replicon identity is conceptualized. Current classification strategies typically rely on individual molecular features, including replication systems, nucleotide composition, conserved markers or sequence similarity^13,26–28,40,41^. However, IRs demonstrate that these features do not always evolve as a coordinated package. They retain extensive chromosomal sequence ancestry while exhibiting plasmid-like compositional characteristics, relaxed evolutionary constraints and distinct replication dynamics, yet lack canonical replication systems associated with either chromosomes or plasmids. No single molecular feature uniquely defines this state; instead, IRs emerge only when multiple independent dimensions are considered together. Consistent with this view, machine-learning models trained on these multidimensional features can distinguish IRs from canonical replicon classes (Fig. 3L; Supplementary Fig. S17). Thus, replicon identity is better understood as an emergent molecular state arising from the integration of diverse evolutionary constraints, rather than as a fixed category defined by a single diagnostic signature.

Our findings also provide a revised perspective on bacterial genome evolution. IRs are preferentially associated with mobilome-enriched chromosomal remodeling hotspots while occupying a distinct topological neighborhood within the mobile plasmid network. These observations indicate that chromosomes and plasmids might be evolutionarily connected through recurrent sequence exchange and genome remodeling. Importantly, our data do not establish the directionality of these events. IRs may arise through chromosomal excision followed by autonomous maintenance, through progressive domestication of mobile plasmids, or through repeated bidirectional exchange between both processes^21–24^. These scenarios are not mutually exclusive and may operate simultaneously across different bacterial lineages or evolutionary timescales. Rather than supporting a single evolutionary pathway, our results suggest that IRs represent a stable molecular state repeatedly revisited during bacterial genome evolution.

Several important questions remain unresolved. Most notably, the molecular mechanisms responsible for replication and segregation of IRs remain unknown, particularly given their apparent lack of recognizable chromosomal or plasmid replication initiators^43–45^. Likewise, comparative genomics alone cannot distinguish whether individual IRs predominantly arise through chromosomal excision, plasmid domestication or repeated transitions between both processes. Addressing these questions will require experimental reconstruction of IR dynamics, direct measurements of replication and segregation, and long-term evolution studies^66,67^.

More broadly, this work shifts the study of bacterial replicons from a binary classification problem toward a quantitative understanding of genomic organization. Rather than asking whether an autonomous replicon is a chromosome or a plasmid, future studies can investigate which molecular state it occupies, how these states are maintained, and how transitions among them contribute to the evolution of bacterial genomes^19,68^.

## Methods

### Calculation of plasmidness

All genomic files were downloaded from the NCBI RefSeq database (GBFF format; accessed on 8 August 2025), restricting to fully sequenced and annotated genomes. Each replicon was classified as either chromosome or plasmid based on RefSeq annotations^29^. Subsequently, DNA sequences belonging to the same genus (the 21 genera with the largest numbers of available sequenced genomes) were pooled into a single FASTA file per genus, and the makeblastdb tool from the NCBI BLAST+ suite (version 2.17.0)^30^ was utilized to construct local BLAST databases.

Each replicon was then used as the query in blastn searches against the corresponding genus-specific BLAST database (parameters: evalue = 1e−50, max_target_seqs = 100,000, max_hsps = 3,000, identity threshold ≥ 90%, outfmt = 6)^30^. For each nucleotide position in a query sequence, we counted the number of aligned plasmid-derived hits (*n_p_*) and chromosome-derived hits (*n_c_*) covering that position. A positional plasmid fraction was defined as 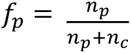. For positions with no alignments, *f_p_* was set to 1 for plasmid queries and 0 for chromosome queries. The plasmidness of a sequence was defined as the mean of *f_p_* across all nucleotide positions.

### Classification of replicons

Replicons were classified based on a combination of the plasmidness metric and sequence length. We first defined typical plasmids as replicons with plasmidness ≥ 0.3. For replicons with plasmidness < 0.3, sizes are first log_10_-transformed and sorted in descending order. We search for the first pair of adjacent entries where the difference in log_10_-size exceeds 0.25. If the raw size of the larger entry in this pair is greater than 1 Mb, the mean raw size of these two entries is taken as the size threshold. If this criterion is not met, replicons are restricted to the size range 100 kb ≤ size ≤ 1 Mb. Within this range, replicons are sorted by log_10_-size in descending order, and the position with the maximum absolute difference between adjacent log_10_-size values is identified. The average length of the two replicons flanking this maximum-jump position is returned as size threshold. If none of the above conditions are satisfied, size threshold is defined as half the minimum replicon size in the filtered dataset. Replicons below this threshold were classified as IRs, whereas those above were classified as chromosomes.

To further assess replicon classification robustness, we performed hierarchical clustering using the two features: size and plasmidness. Hierarchical clustering was performed using the average-linkage method^29^, with normalized plasmidness and normalized log_10_-transformed replicon size as input features. Merge-distance values from the latter-stage merging events of the clustering tree were extracted, and their first- and second-order differences were calculated. The cutting threshold for tree partitioning was determined based on the maximum absolute value of the second-order difference; the final cutoff was the mean of three consecutive merge distances around this position. The dendrogram was partitioned under the distance criterion to assign cluster membership for each sample.

Bootstrap resampling was applied to assess the robustness of the inferred cluster number. Resampled datasets were generated by drawing samples from the original dataset with replacement. The full hierarchical-clustering and automatic threshold-partitioning workflow was repeated on each bootstrap replicate to obtain cluster counts. We counted how many bootstrap replicates reproduced the same cluster number as observed in the original dataset and calculated its fraction among all replicates as a robustness metric.

We further performed chromosome-plasmid classification for plasmids and IRs using four independent tools: PlasFlow^27^, Plasmer^40^, RFPlasmid^28^, and Deeplasmid^41^, using default or recommended parameters. We additionally applied chromid-finder^42^ to all replicons, together with chromids identified or predicted in previously published literature^14,42^, to evaluate the presence of chromid-like replicons.

### Genome re-assembly

To assess whether the IRs represent genome assembly artifacts, we performed genome re-assembly for a subset of *Escherichia* (family Enterobacteriaceae) genomes with publicly available short-read and long-read sequencing data. Genomes were selected from the SRA^69^ based on the availability of paired Illumina short reads and long-read datasets. Hybrid genome assemblies were generated using three independent assemblers, Unicycler (v0.5.1)^32^, Hybracter (v0.13.0)^33^, and Flye (v2.9.6)^34^, under default or recommended settings for assembly. For each assembly, contigs ≥1 kb were retained for downstream analysis. The plasmidness metric was then computed for each contig using the pipeline described above.

### Replication origin prediction and protein annotation

Replication origin (*oriC*) sites were predicted for all replicons using ORCA^44^. Plasmid replication origins (*oriV*) were further identified for typical plasmids and IRs using OriV-Finder^45^ under default parameters. Protein-coding sequences were annotated using eggNOG-mapper (v2.1.13)^70^ with bacterial taxonomic scope and DIAMOND-based alignment^71^, to retrieve gene names, COG identifiers^51^ and COG functional categories for each sequence. Protein percentage for each COG category was computed on the basis of protein counts. For each replicon group, the numerator was the count of proteins annotated to the focal COG category, while the denominator represented the total number of proteins with valid amino-acid translations within the same replicon. The frequencies of COG categories were calculated for each replicon, which were then used for principal-component analysis (PCA)^52^.

### Genomic composition and *k*-mer analysis

GC content and *k*-mer frequencies were computed across both forward and reverse complement strands (*k* = 3, 4, 5). To compare GC content across different replicon types, linear fitting was performed using the GC content of each replicon as the y-axis value and the GC content of its corresponding whole-genome as the x-axis value. The coefficient of determination (*R*²), slope and intercept were calculated accordingly.

For individual genera, 200 samples were randomly sampled from each replicon class (all available samples were used when fewer than 200 were present). PCA^52^ was conducted on *k*-mer frequencies for different *k* values, and ANOSIM *R* values^56^ were computed between replicon classes. This replicon-sampling procedure was repeated five times.

### Chromosomal fragment controls

To control for length-dependent compositional biases, chromosomal fragments were generated by randomly sampling chromosomal segments matching the length distribution of plasmid and IRs. In each sampling iteration, one sequence was randomly picked from the combined set of plasmids and IRs, and only its length was retained as the target fragment length. A complete chromosome was then randomly selected from the chromosome pool, from which a chromosomal segment of this target length was randomly excised. When the target length exceeded the total size of the sampled chromosome, the full-length chromosome was kept as the resulting fragment.

### Pseudogene and copy number quantification

Pseudogenes were defined as coding sequence (CDS) annotations lacking a translated protein sequence in GBFF records^57^. The pseudogene proportion was calculated as the number of pseudogene CDS annotations divided by the total number of all CDS annotations retrieved from GBFF records. Replicon copy-number information was retrieved from published datasets^58^ and assigned to contigs using sequence identifiers.

### Calculation of embedding vectors from foundation model

Embedding-vector computation was performed only for *Escherichia* strains. First, up to 200 sequences were randomly sampled for each replicon category; all available sequences were used when fewer than 200 samples existed. DNA sequences exceeding the length threshold were randomly cropped into sub-fragments, whereas sequences shorter than the threshold were kept in full length. Sequences were fed into pre-trained DNA foundation models, and hidden-state outputs were obtained with gradient computation disabled. One-dimensional sequence embeddings were generated via mean pooling over the token dimension, which were subsequently used for PCA^52^.

Three pre-trained DNA foundation models with different parameter scales were adopted for embedding extraction, including Evo2_1b_base.pt^59^, gLM2_650M^60^, and nucleotide-transformer-2.5b-multi-species^61^, with distinct sequence cropping and feature aggregation strategies for each model. Specifically, the Evo2 model cropped input sequences to a maximum length of 30,000 bp, extracted hidden features from the *blocks.24.mlp.l3* layer, and generated sequence embeddings by mean pooling across all tokens. The gLM2 model adopted a longer cropping threshold of 50,000 bp and obtained final embeddings via mean pooling over all tokens from the last hidden state layer. For the nucleotide-transformer model, raw DNA sequences were first physically truncated to 6,000 bp, followed by tokenizer-based truncation with a maximum length of 1000 tokens; the redundant special learnable token was discarded, and the final sequence embedding was calculated by averaging only the valid intermediate tokens.

### Machine learning classification of replicon categories

Classifiers were constructed using machine learning algorithms to distinguish the three types of replicons based on genomic compositional features. The input features included *k*-mer frequencies (*k* = 3, 4, 5) and COG category frequencies. Two comparative experimental settings were designed, with or without the incorporation of sequence size features. For each setting, corresponding *k*-mer features, COG frequency features, and optional sequence size features were integrated to construct the feature matrix.

All samples were randomly and stratified split into training and test datasets at a ratio of 7:3. Three classic machine learning classifiers were applied, including multinomial Logistic Regression, Random Forest^72^, and linear Support Vector Machine (Linear SVM)^73^. Feature standardization was performed prior to the training of linear models. Balanced class weights were adopted for all models to reduce classification bias caused by imbalanced sample sizes among different replicon groups.

The predictive performance of each model was comprehensively evaluated on the test set using overall accuracy, class-specific precision, recall, F1-score, and confusion matrix. A one-versus-rest strategy was applied to quantify multi-classification performance. The receiver-operating-characteristic (ROC) and precision-recall curve data corresponding to each replicon category were recorded, and the area under the ROC curve (AUROC) and area under the precision-recall curve (AUPRC) values were calculated accordingly.

### Construction of chromosome clustering trees

To quantify genome-wide sequence similarity, we computed pairwise alignment-based similarity profiles between all replicons using aggregated BLAST bitscores (as defined in the plasmidness pipeline)^30^. For genomes containing multiple chromosomal fragments, chromosomal signals were merged by summing bitscore contributions across different chromosomes and normalizing by total chromosomal length, yielding a length-adjusted similarity representation.

These normalized bitscore profiles were assembled into a genome-level feature matrix. Pairwise distances between genomes were computed using Euclidean distance, and hierarchical clustering was performed using the average-linkage method^31^. The resulting dendrograms were converted into phylogenetic tree formats for visualization.

Statistical robustness for clustering structure was assessed using bootstrap resampling. In each iteration, the feature matrix was resampled with replacement, and clustering was recomputed using the same pipeline. Branch support values were defined as the proportion of bootstrap trees in which each clade was recovered across 1,000 replicates.

### Associations between replicons

To quantify sequence-level associations between replicons, we defined an alignment-based overlap criterion using BLAST-derived coverage intervals. A replicon was considered associated with a chromosome if alignments between the replicon and the chromosome covered at least two-thirds of the replicon length. For associations between non-chromosomal replicons (i.e., IRs and plasmids), an edge was defined when the aligned regions covered at least two-thirds of either replicon, ensuring sensitivity to asymmetric length distributions. Based on these criteria, we constructed a replicon association network in which nodes represent individual replicons and edges represent significant sequence overlap. Network visualization was performed using Cytoscape (v3.10.4)^74^.

For each IR longer than 10 kb, BLAST alignments against chromosomes with sequence identity ≥90 % were filtered and sorted in descending order of bitscore. Only the best alignment per chromosome was retained, and the top-five chromosomes were selected. The start and end coordinates of the aligned segment on the chromosome were extracted, and each alignment boundary was extended by 2500 bp both upstream and downstream to obtain the upstream and downstream flanking regions surrounding the putative fission-fusion sites of IRs on the chromosome. All CDS features within these flanking intervals were extracted, and their corresponding COG annotation information was recorded.

A random background control was implemented: a random segment of the same length as the real aligned fragment was sampled from the identical chromosome. This random segment was also extended by 2500 bp on both sides, and CDS features together with COG annotations within this interval were collected to serve as the random background.

### Mobility and antimicrobial resistance traits of replicons

Mobility and antimicrobial resistance (AMR) traits were annotated for typical plasmids and IRs. Replicon mobility was predicted using MOB-typer (v3.1.9)^75^ with default parameters, classifying replicons based on established mobility marker genes. AMR genes were identified using AMRFinderPlus (v4.2.5)^76^ with the 2025-12-03 database release. A replicon was defined as an AMR-positive replicon if it contained at least one confidently annotated AMR gene.

### Analysis of plasmids assembled from metagenomic data

Plasmid sequences were retrieved from the IMG/PR database^65^. Only plasmids assembled from metagenomic data were retained. We filtered plasmids belonging to the 21 bacterial genera selected in this study and calculated their plasmidness scores using the aforementioned pipeline. These plasmids were classified into replicon categories on a per-genus basis and further categorized according to ecosystem metadata from the database. MOB-typer^75^ and AMRFinderPlus^76^ were used to identify mobility features and AMR determinants.

## Data Availability

All the data associated with this work are available at the GitHub repository (https://github.com/Zpresitong/Plasmidness.git).

## Code Availability

All codes are available at the GitHub repository (https://github.com/Zpresitong/Plasmidness.git).

## Acknowledgement

This study was supported by the National Key R&D Program of China (2024YFA0920200 to TW), the National Natural Science Foundation of China (12401660 and 32470701 to TW), and the Shenzhen Institute of Synthetic Biology Scientific Research Program (HSE499011086 to TW). We are grateful to the Shenzhen Infrastructure for Synthetic Biology for providing instrument support and technical assistance.

