## Supplemental File # 1 for "Hidden molecular states of bacterial replicons beyond the chromosome–plasmid dichotomy"

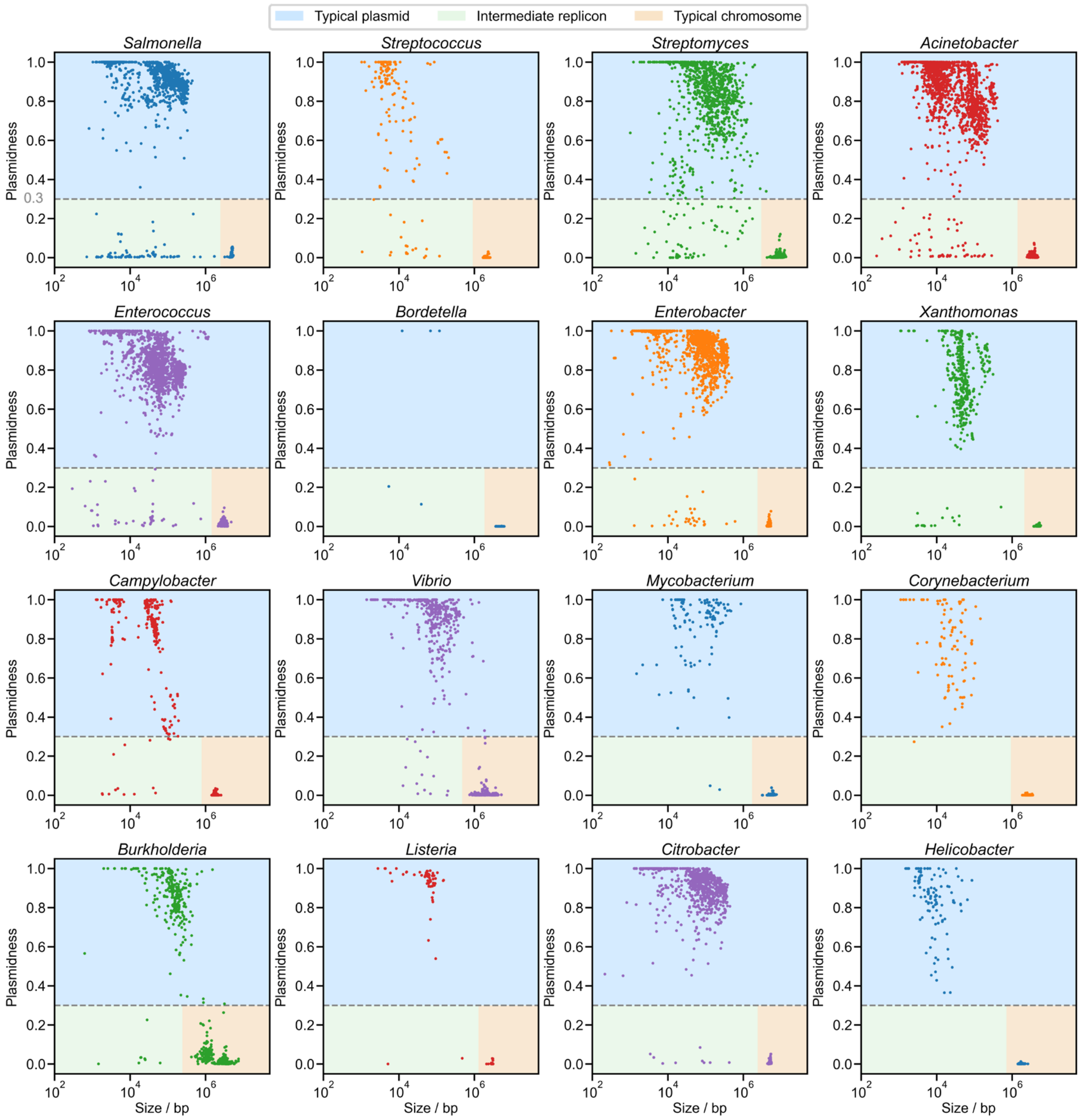


**Supplementary Fig. S1 | Conservation of the three-state replicon organization across bacterial genera.** Replicon landscapes from 16 representative bacterial genera reveal the recurrent emergence of three molecular states corresponding to typical plasmids, intermediate replicons (IRs), and chromosomes.

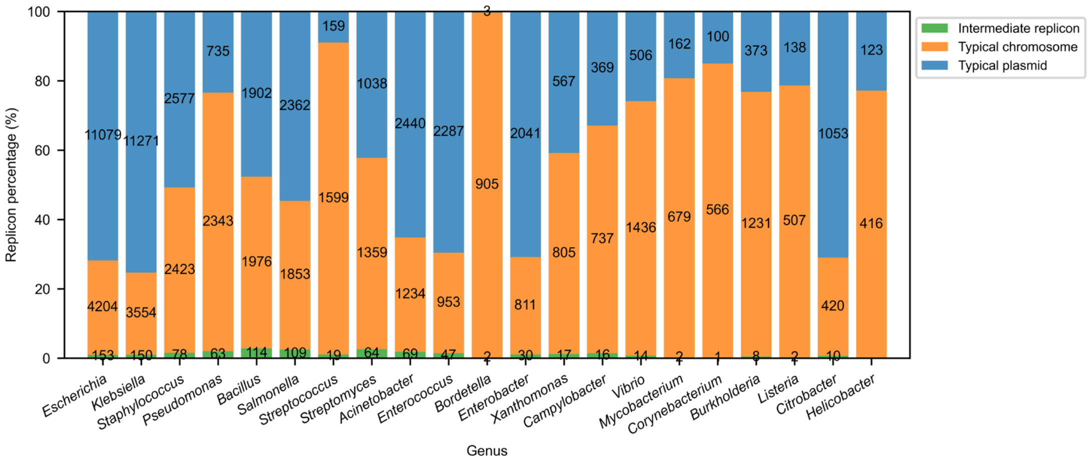


**Supplementary Fig. S2 | Relative abundance of replicon states across bacterial genera.** The proportion of intermediate replicons (IRs) varies across bacterial genera but remains a minor component of the total replicon population, accounting for less than 3% in all examined genera.

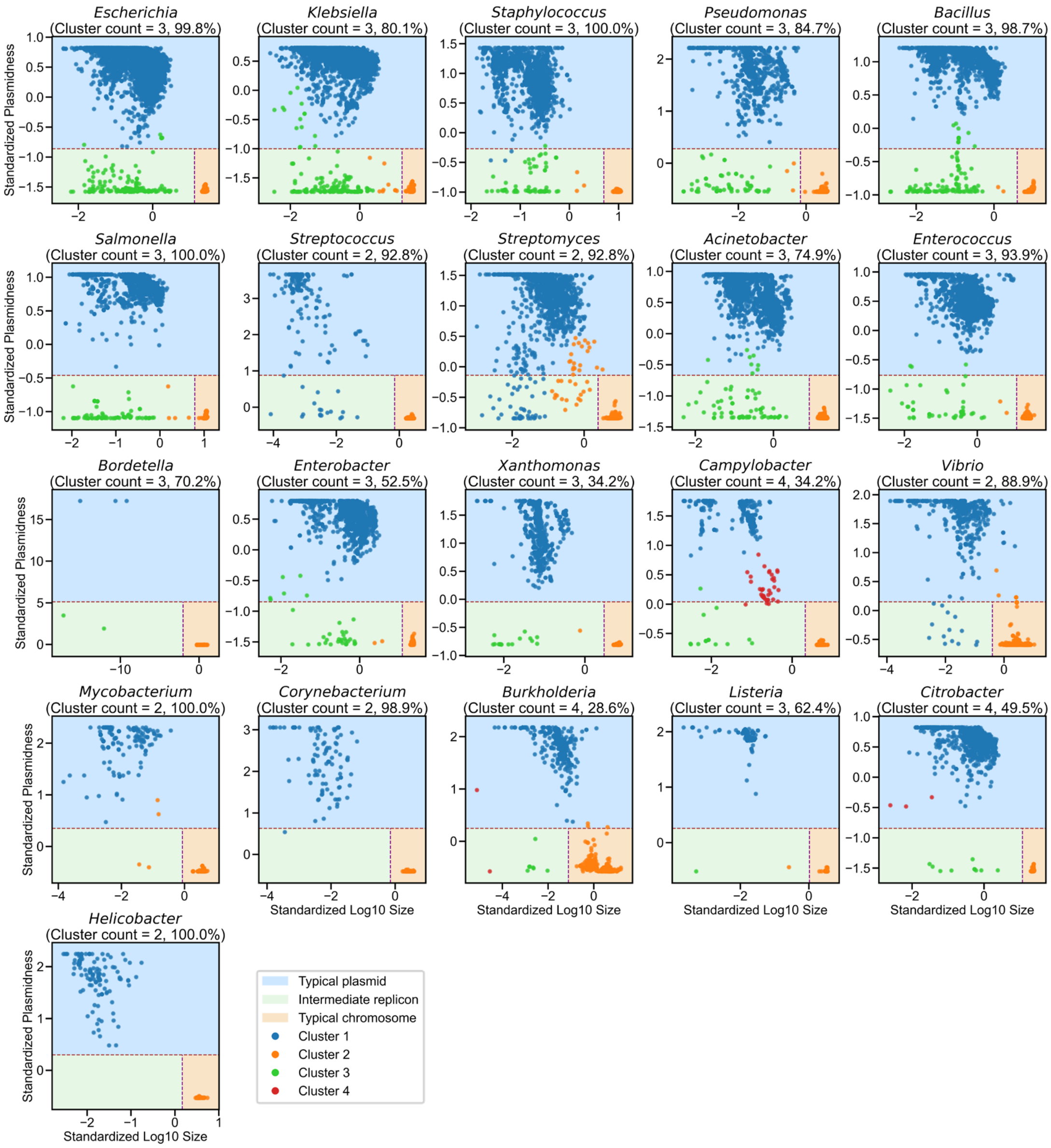


**Supplementary Fig. S3 | Unsupervised validation of the three-state replicon organization.** Replicon plasmidness and log10-transformed size were standardized before average-linkage hierarchical clustering. Independent clustering of these two quantitative features recapitulated the three replicon groups identified in this study. Bootstrap resampling was used to evaluate clustering stability; percentages indicate the fraction of resampled datasets recovering the corresponding number of clusters. Colored regions denote the predefined replicon states, whereas colored points represent clusters identified by hierarchical clustering.

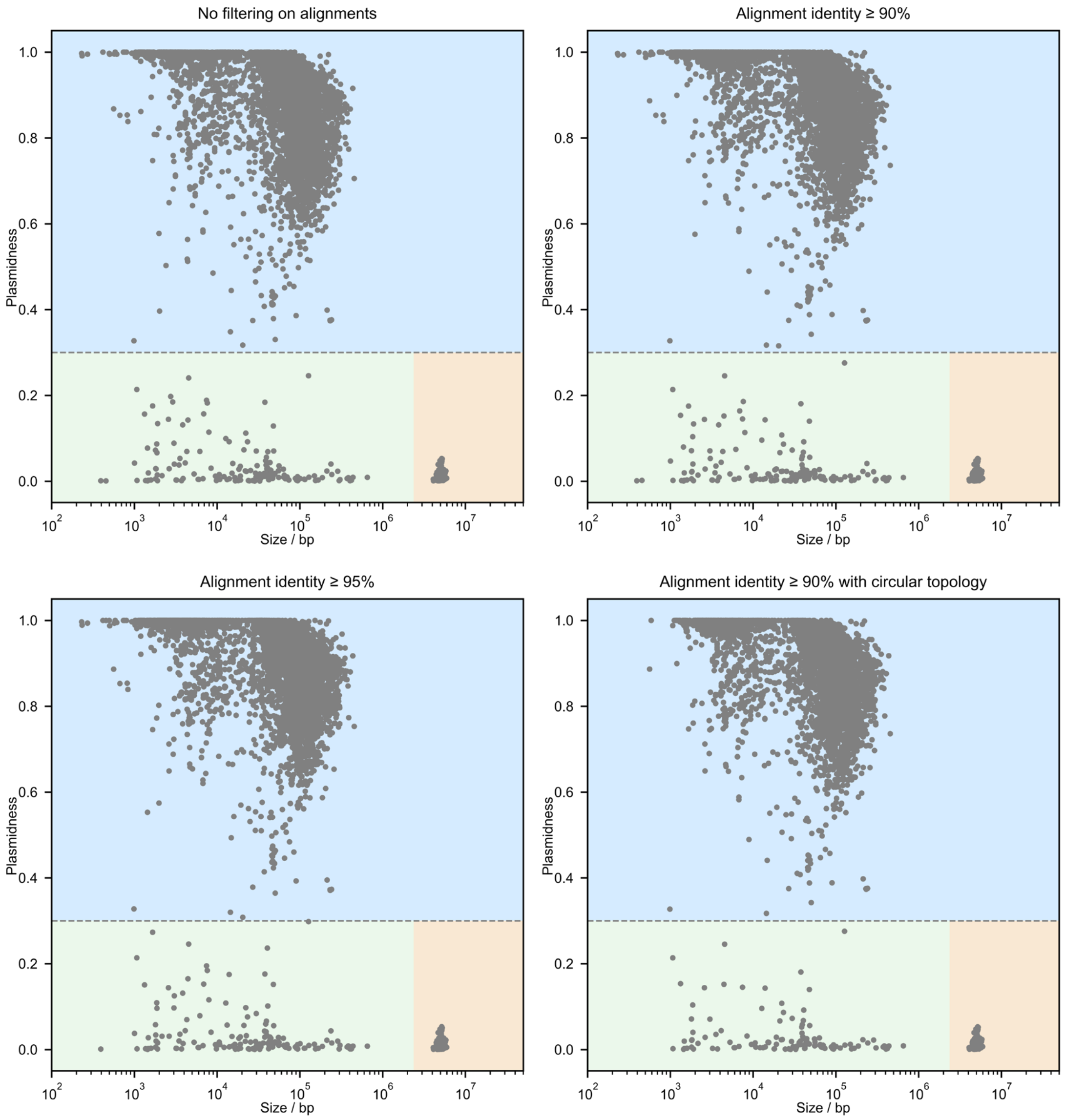


**Supplementary Fig. S4 | Robustness of the three-state replicon organization to sequence filtering criteria.** Replicon distributions in *Escherichia* remain consistently partitioned into three groups across alternative alignment identity thresholds and after restricting analyses to circularly assembled replicons.

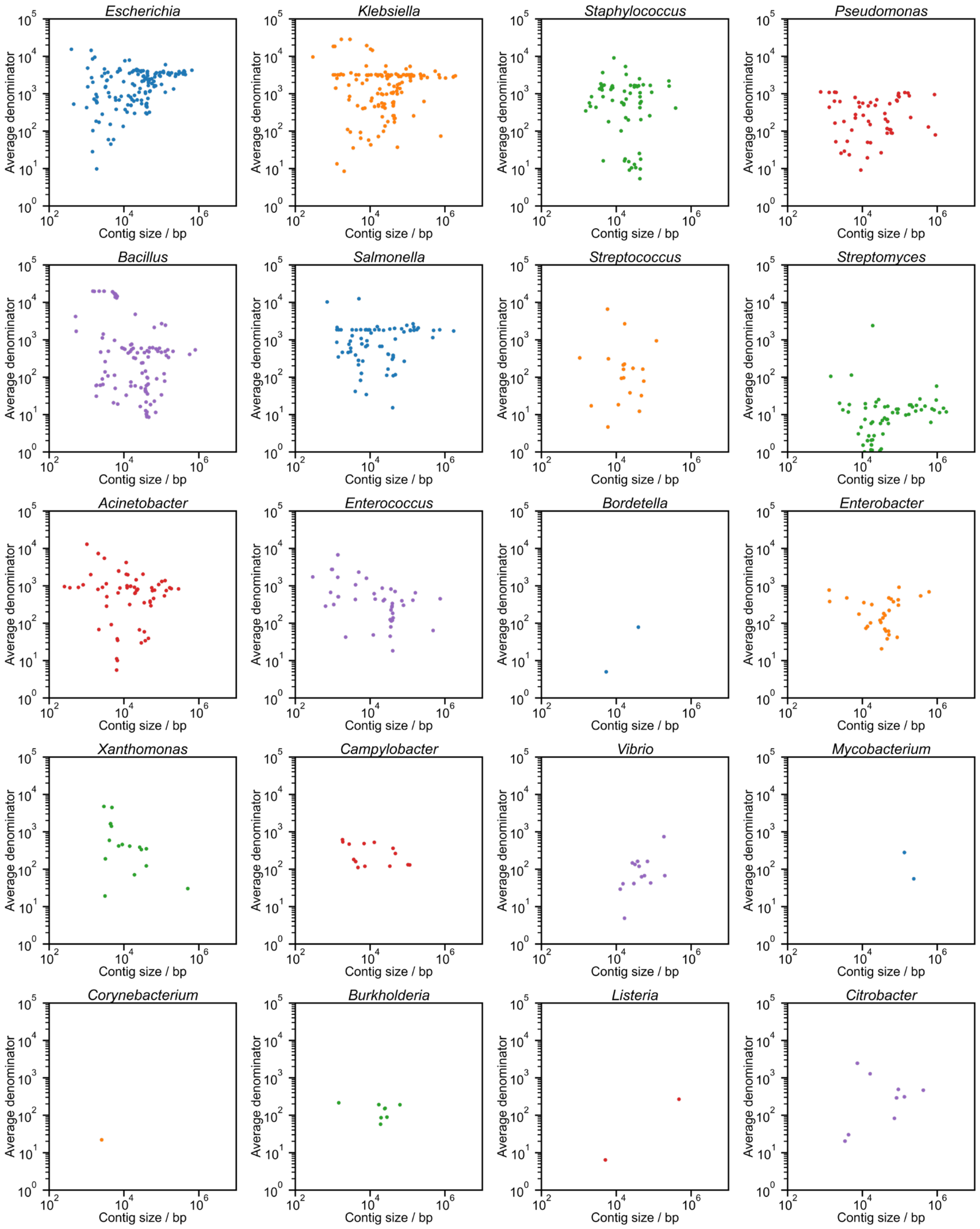


**Supplementary Fig. S5 | Alignment depth underlying replicon plasmidness estimation.** The average number of aligned sequences contributing to each nucleotide position of intermediate replicons is approximately 1,000 across most bacterial genera, with the majority of replicons supported by at least 100 aligned sequences per site.

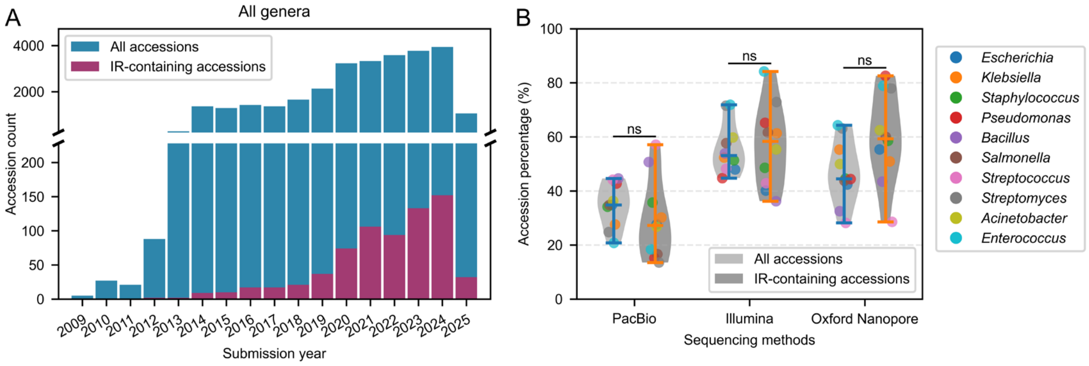


**Supplementary Fig. S6 | Assessment of temporal and sequencing biases in IR-containing genomes.**

(A) Submission-year distributions of IR-containing genome accessions against all genome accessions.

(B) Accession proportion stratified by sequencing-technology categories. The y-axis denotes accession proportion, representing the fraction of genomes using a given sequencing method relative to all accessions within each category bin. Two-sided Welch's *t*-test was used for group comparisons. Significance markers: ***, $P < 0.001$; **, $P < 0.01$; *, $P < 0.05$; ns, non-significant ($P \geq 0.05$).

Comparable distributions of IR-containing genomes versus the total genome set across submission years and sequencing platforms indicate that IR identification is not driven by temporal sampling bias or sequencing-technology effects.

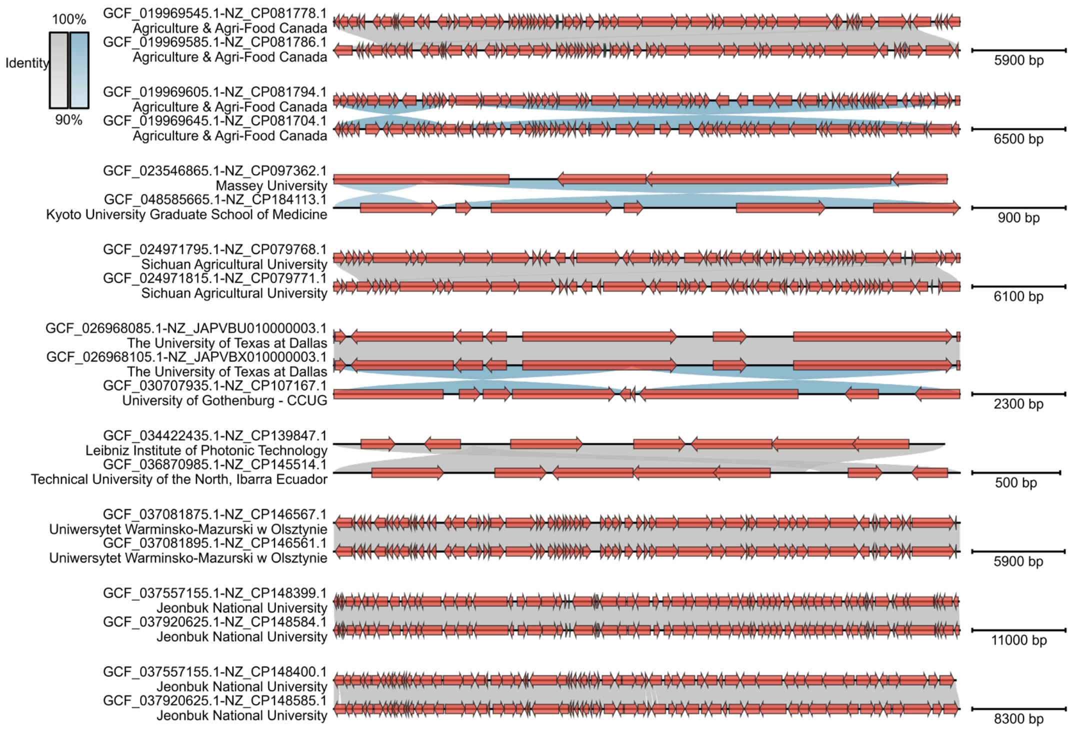


**Supplementary Fig. S7 | Recurrence of highly similar intermediate replicons across independent *Escherichia* genomes.** Highly similar or identical IR sequences are detected across distinct *Escherichia* genomes. Red arrows indicate coding sequences (CDSs), with sequence accession numbers and submitting institutions shown on the left.

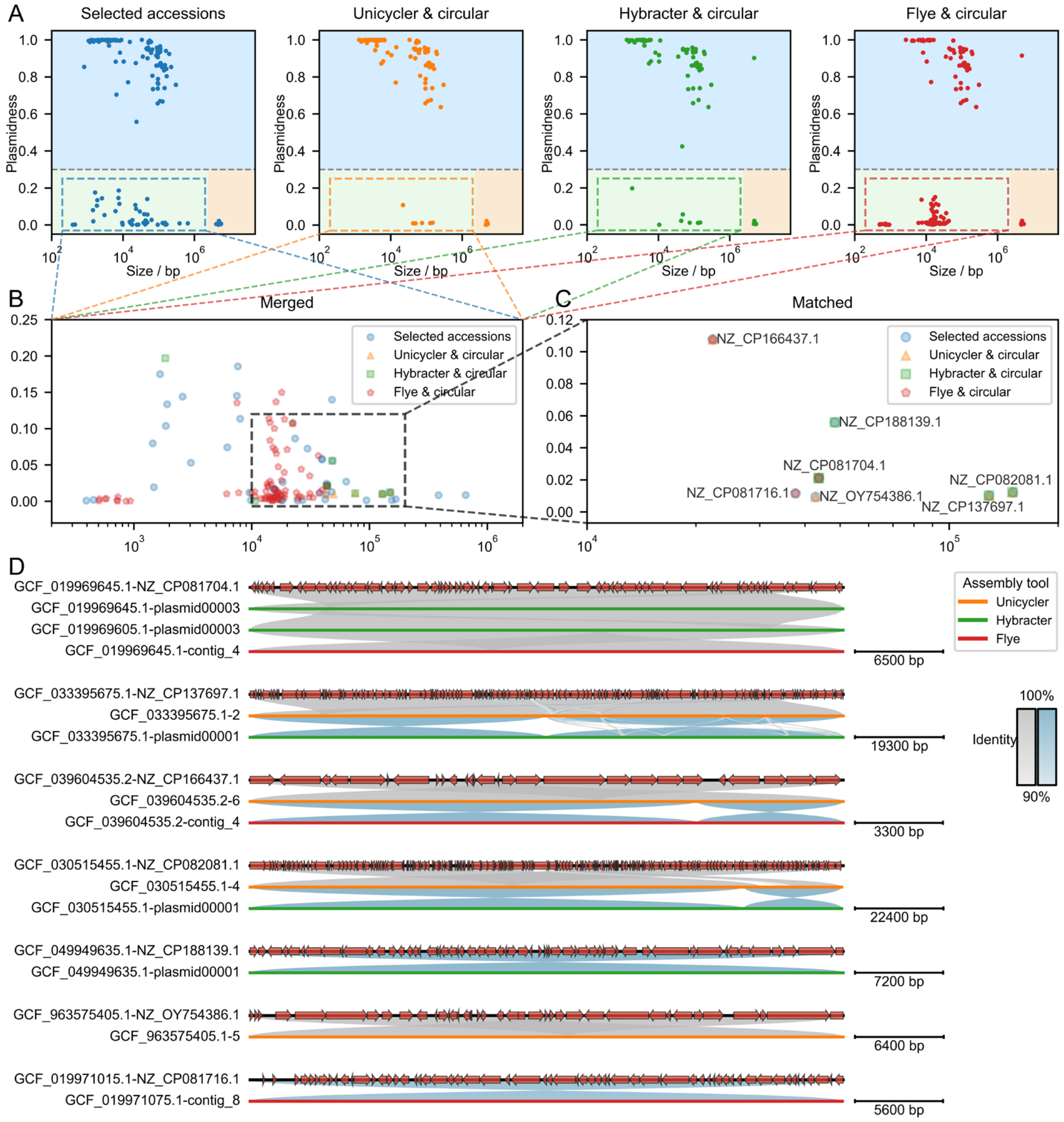


**Supplementary Fig. S8 | Robust recovery of intermediate replicons across independent genome assemblies.**

(A) Size-plasmidness distribution of replicons from *Escherichia* accessions selected for reassembly, together with outputs from different assembly tools (only circularized assemblies were retained).

(B) Merged results of the intermediate-replicon blocks derived from the four scatter plots.

(C) Assembly outputs screened across three assembly methods whose size-plasmidness properties match those of intermediate replicons in the original accessions.

(D) Sequence-alignment results between screened assembly outputs and original intermediate replicons.

Independent reassembly of representative genomes using alternative assembly pipelines consistently recovers circular intermediate replicons, supporting their status as genuine genomic entities rather than assembly artifacts.

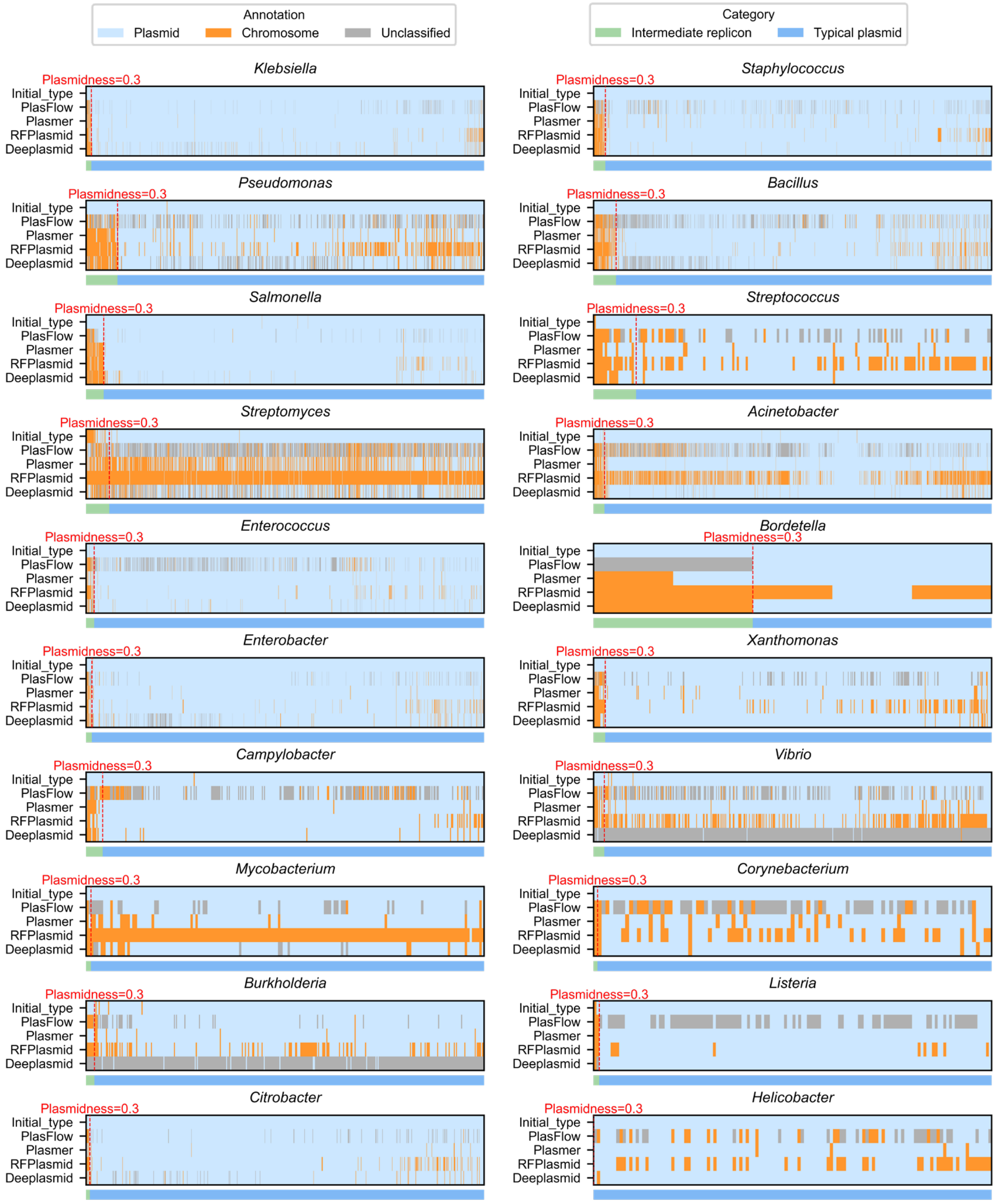


**Supplementary Fig. S9 | Classification ambiguity of intermediate replicons across bacterial genera.** Four independent replicon classification methods consistently assign typical plasmids to the plasmid category, whereas intermediate replicons show variable assignments, including frequent classification as chromosomes. Classification patterns are consistent across bacterial genera, with method-specific differences reflecting the distinct molecular characteristics of intermediate replicons.

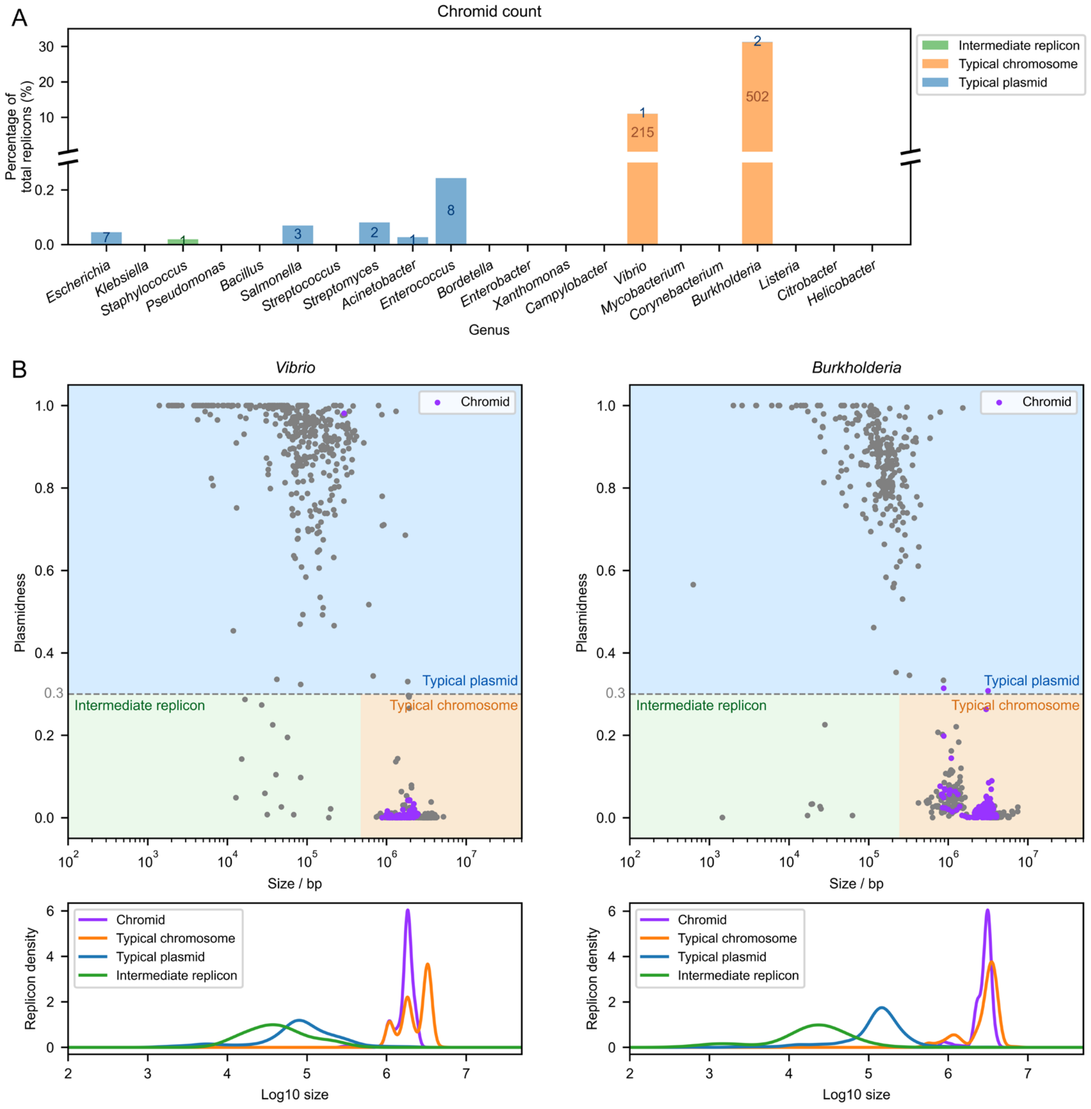


**Supplementary Fig. S10 | Screening results for chromid-class replicons.**

(A) Results of chromid screening for replicons across all bacterial genera. Color blocks indicate the replicon category from which each chromid was identified; values represent replicon counts.

(B) Chromid-screening distributions for the two genera (*Vibrio* and *Burkholderia*) with the largest number of chromids. Shown are the size-plasmidness distribution of chromids and replicon-density curves of different replicon types against $\log_{10}$-transformed size.

Chromid screening across all replicons reveals minimal overlap between intermediate replicons and chromids. Most detected chromids are restricted to *Vibrio* and *Burkholderia* lineages and occupy the chromosomal region of the replicon landscape.

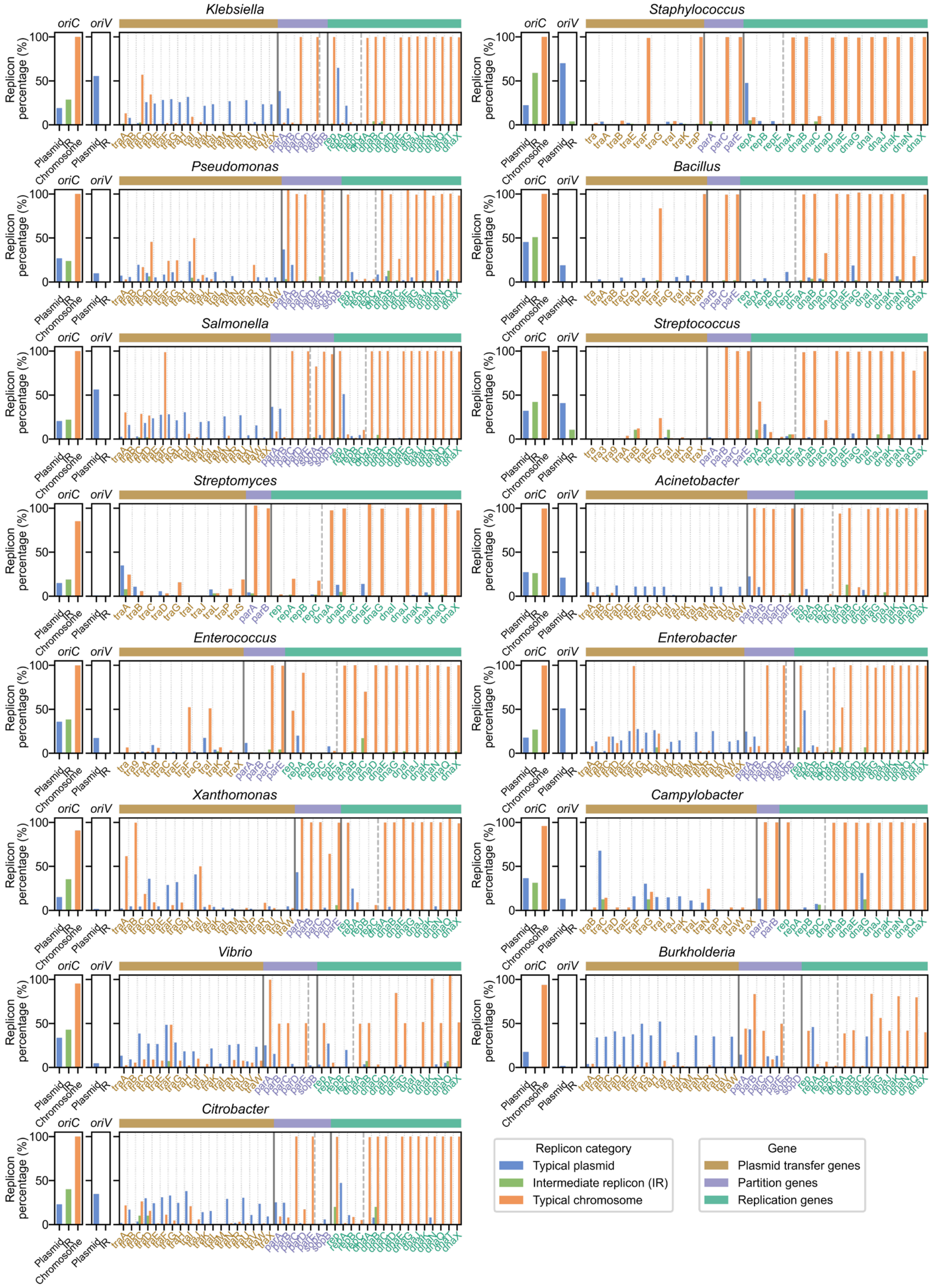


**Supplementary Fig. S11 | Replication-associated features and functional core-gene markers of replicons across bacterial genera.** Replication origin prediction reveal that intermediate replicons rarely contain canonical *oriV* elements, while a subset harbors *oriC*-associated regions. Their functional core-gene repertoires differ from those of typical plasmids and chromosomes across bacterial genera.

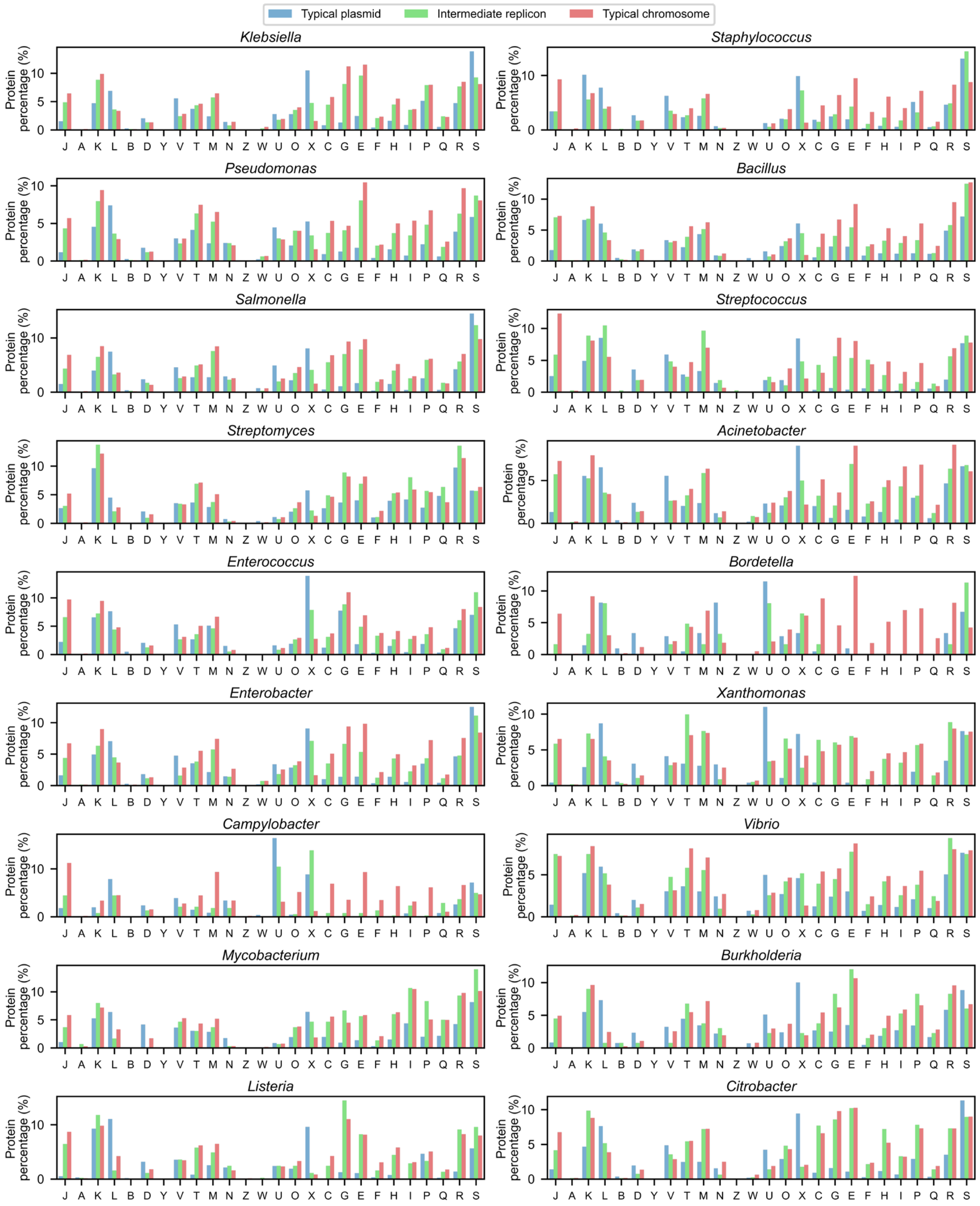


**Supplementary Fig. S12 | Functional composition of replicon states based on COG categories across bacterial genera.**

The relative abundance of COG functional categories in intermediate replicons (IRs) shows an intermediate profile between chromosomes and typical plasmids across bacterial genera. The y-axis (protein percentage) represents the proportion of protein counts for each category relative to the total number of amino-acid-containing proteins within the corresponding replicon group. Letter annotations correspond to COG functional categories as defined below.

Information storage and processing: (J) Translation, ribosomal structure and biogenesis; (A) RNA processing and modification; (K) Transcription; (L) Replication, recombination and repair; (B) Chromatin structure and dynamics.

Cellular processes and signaling: (D) Cell cycle control, cell division, chromosome partitioning; (Y)

Nuclear structure; (V) Defense mechanisms; (T) Signal transduction mechanisms; (M) Cell wall/membrane/envelope biogenesis; (N) Cell motility; (Z) Cytoskeleton; (W) Extracellular structures; (U) Intracellular trafficking, secretion, and vesicular transport; (O) Posttranslational modification, protein turnover, chaperones; (X) Mobilome: prophages, transposons.
Metabolism: (C) Energy production and conversion; (G) Carbohydrate transport and metabolism; (E) Amino acid transport and metabolism; (F) Nucleotide transport and metabolism; (H) Coenzyme transport and metabolism; (I) Lipid transport and metabolism; (P) Inorganic ion transport and metabolism; (Q) Secondary metabolites biosynthesis, transport and catabolism.
Poorly characterized: (R) General function prediction only; (S) Function unknown.

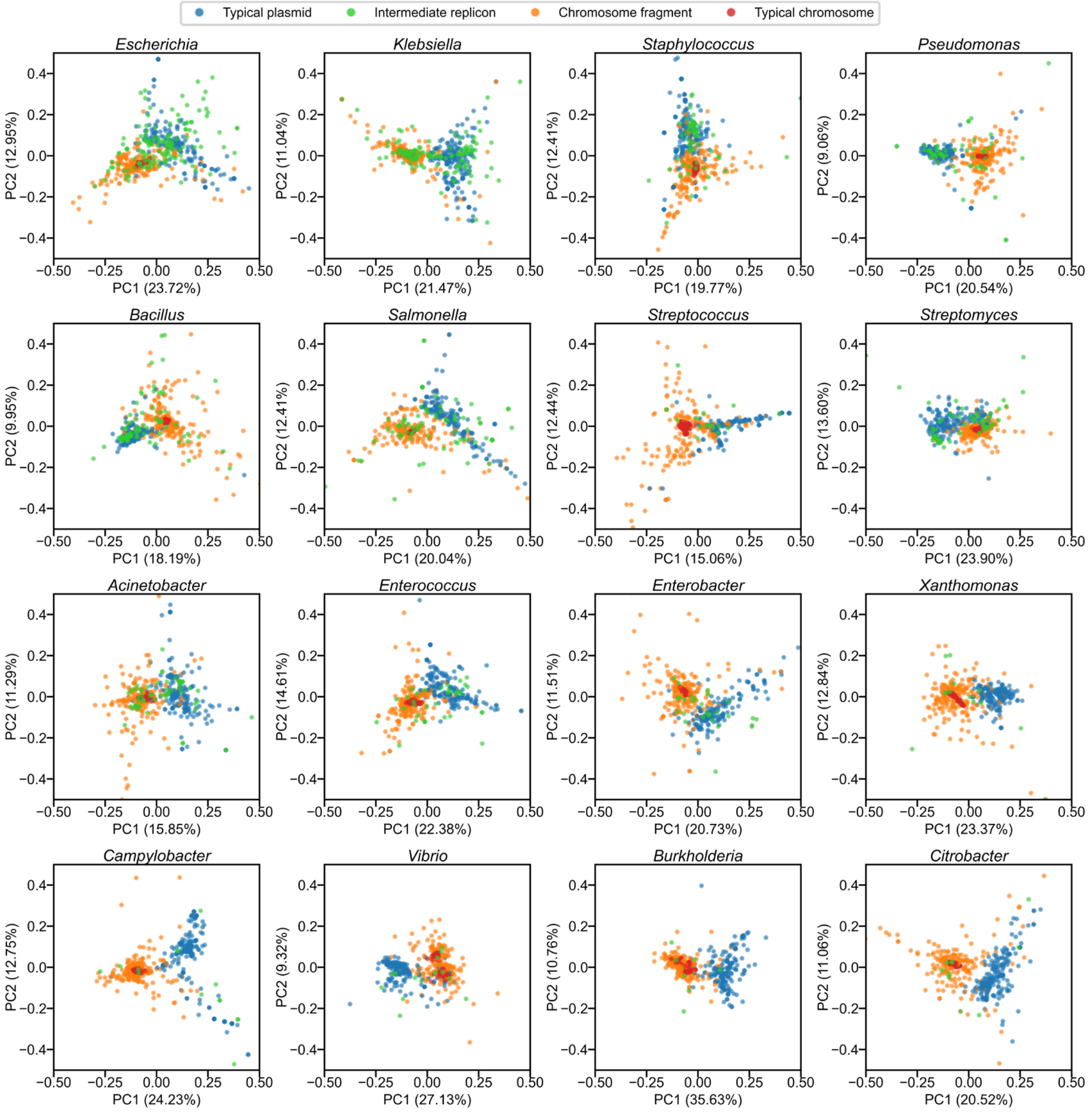


**Supplementary Fig. S13 | Functional landscape of replicon states based on COG-category composition.** Principal-component analysis (PCA) of COG-category frequencies reveals distinct functional distributions of typical plasmids and chromosomal fragments. Intermediate replicons (IRs) occupy an intermediate region of functional space between these two canonical replicon states across bacterial genera containing at least five IRs. PC1 and PC2 denote the first and second principal components; the percentage values on each axis represent the proportion of total variance explained by that principal component.

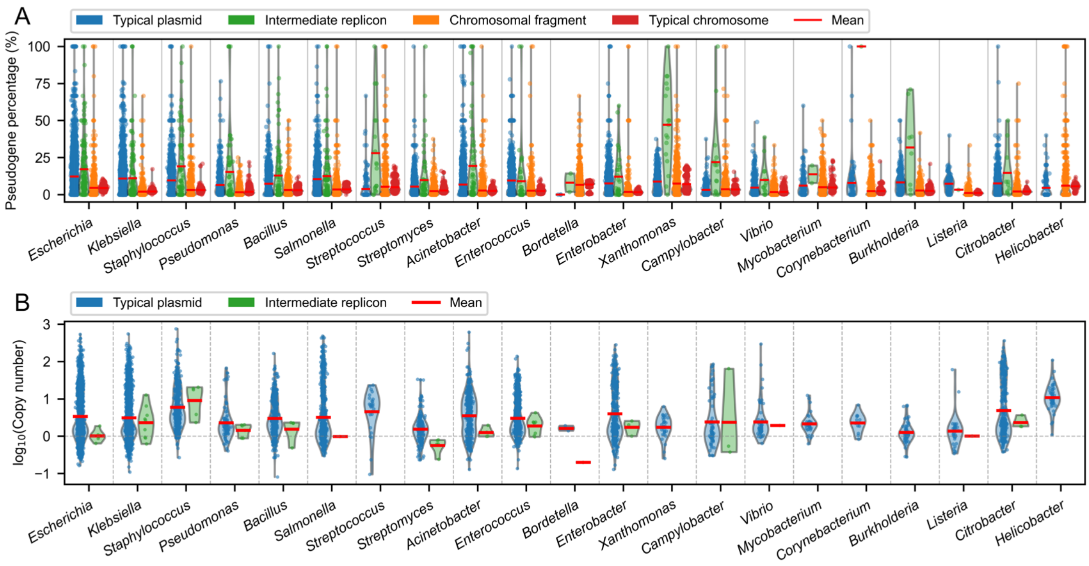


**Supplementary Fig. S14 | Evolutionary constraint and copy numbers of intermediate replicons across bacterial genera.**

(A) Statistics of pseudogene percentage for different sequence types. Pseudogene percentage was calculated for each individual replicon or selected sequence as the fraction of pseudogene CDS among all CDS features from GBFF records.

(B) Copy-number statistics for different replicon types.

Across bacterial genera, intermediate replicons (IRs) exhibit elevated pseudogene frequencies compared with chromosomes, whereas size-matched chromosomal fragments show pseudogene levels comparable to intact chromosomes. IRs also display low copy-numbers relative to typical plasmids, while remaining slightly above chromosomal baseline levels.

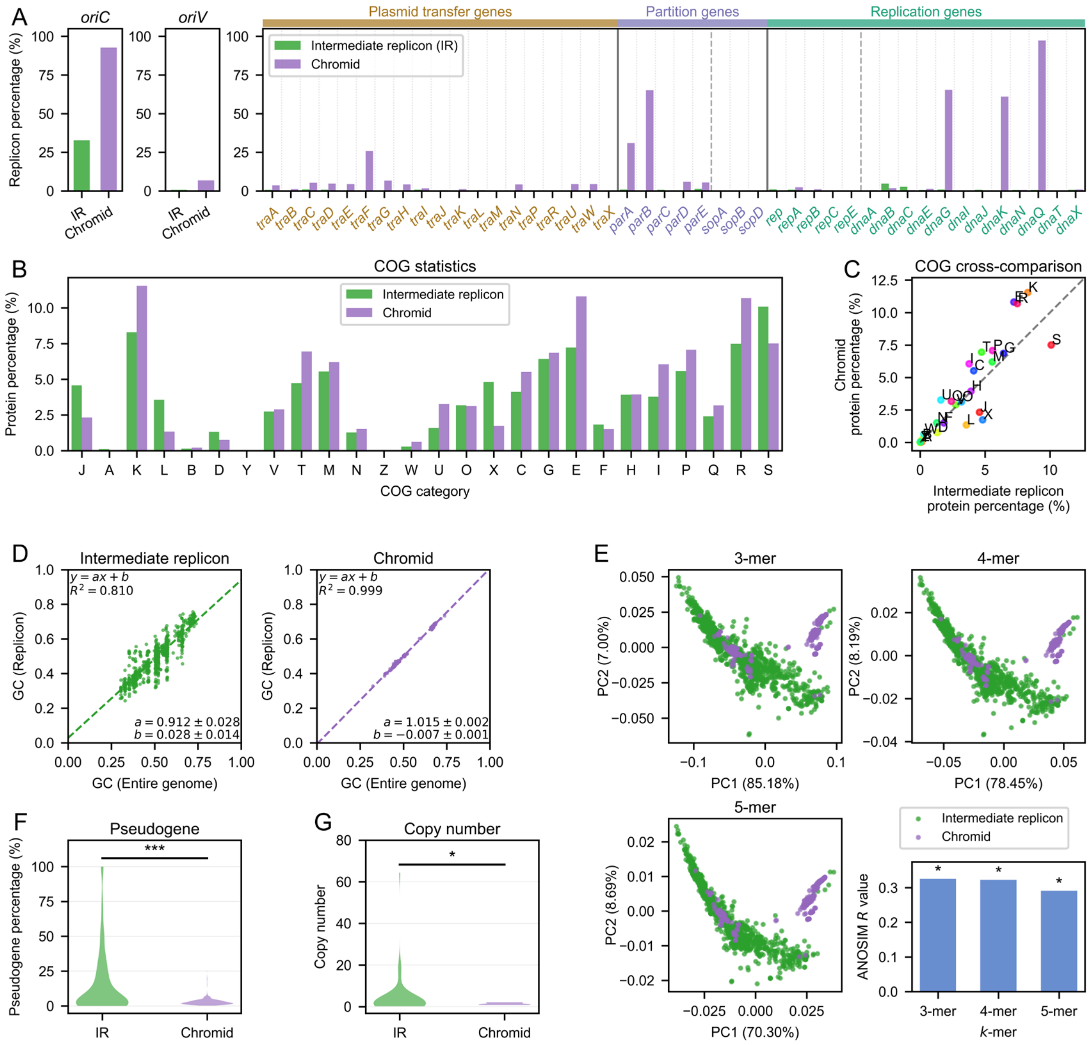


**Supplementary Fig. S15 | Distinct molecular features of intermediate replicons and chromids.**

(A) Statistical comparisons of replication-associated features and functional core-gene markers between IRs and chromids.

(B) Functional composition of IRs versus chromids based on COG categories. Protein percentage represents the proportion of protein counts for each category relative to the total number of proteins.

(C) Comparison of COG proportions between IRs and chromids. The *x*- and *y*-values for each COG category correspond to the statistical outputs of IRs and chromids, respectively.

(D) GC content distributions of IRs and chromids. Uncertainties for the slope and intercept represent the half-widths of the 95% confidence intervals.

(E) Sequence composition landscape and ANOSIM-based similarity comparisons based on *k*-mer profiles. PC1 and PC2 denote the first and second principal components; the percentage values on each axis represent the proportion of total variance explained by that principal component. Significance levels of ANOSIM comparisons: * $p < 0.05$, ** $p < 0.01$, *** $p < 0.001$, ns, not significant ($p \geq 0.05$).

(F) Statistical results of pseudogene percentage. Pseudogene percentage was calculated for each replicon as the fraction of pseudogene CDS among all CDS features from GBFF records.

(G) Statistical results of copy number. Two-sided Welch's t-test was performed for pseudogene proportions and replicon copy number. Significance levels are indicated as: * $P < 0.05$, ** $P < 0.01$,

*** $P < 0.001$; ns, not significant ($P \geq 0.05$).

Intermediate replicons (IRs) differ from chromids across multiple molecular dimensions, including functional composition, sequence characteristics, pseudogene accumulation and copy-number profiles.

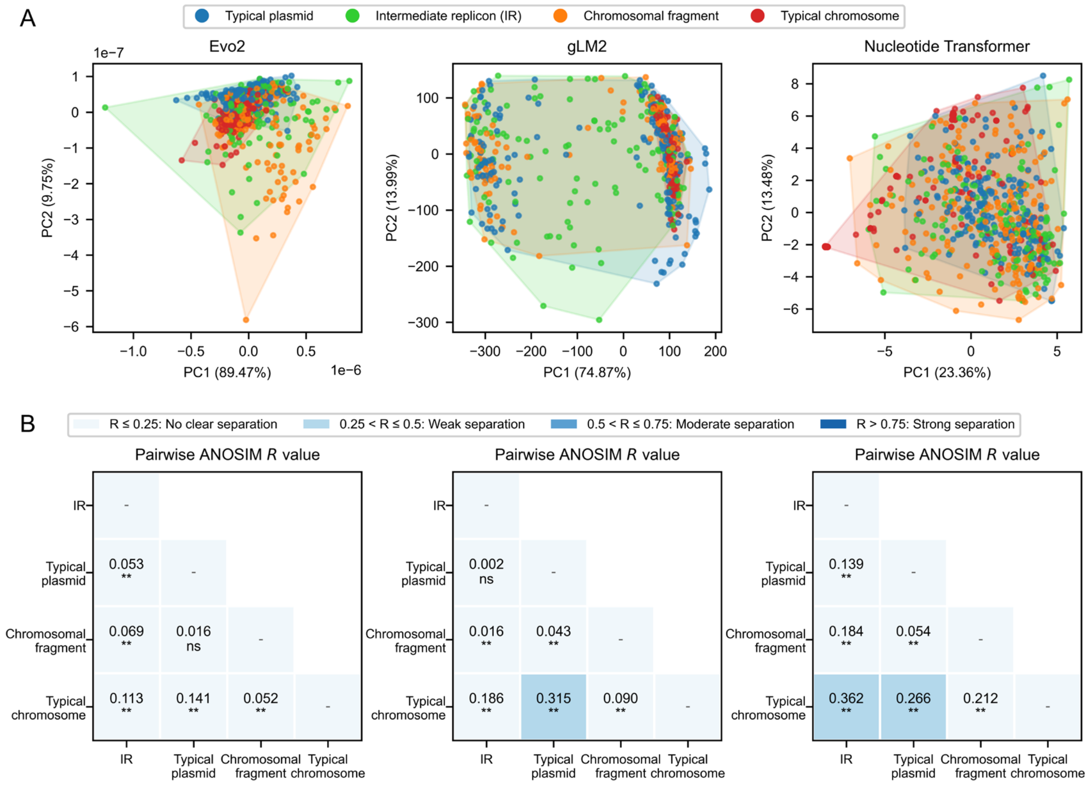


**Supplementary Fig. S16 | Sequence representation of intermediate replicons in genomic foundation models.**

(A) Principal-component analysis of embedding vectors generated by multiple genomic foundation models. The result shows limited separation of intermediate replicons (IRs) from canonical replicon classes.

(B) Pairwise ANOSIM analyses based on embedding vectors. Analyses reveal that IRs exhibit intermediate sequence representation patterns relative to typical plasmids and chromosomal fragments. Asterisks indicate significance levels of ANOSIM comparisons: * $p < 0.05$, ** $p < 0.01$, *** $p < 0.001$, ns, not significant ($p \geq 0.05$).

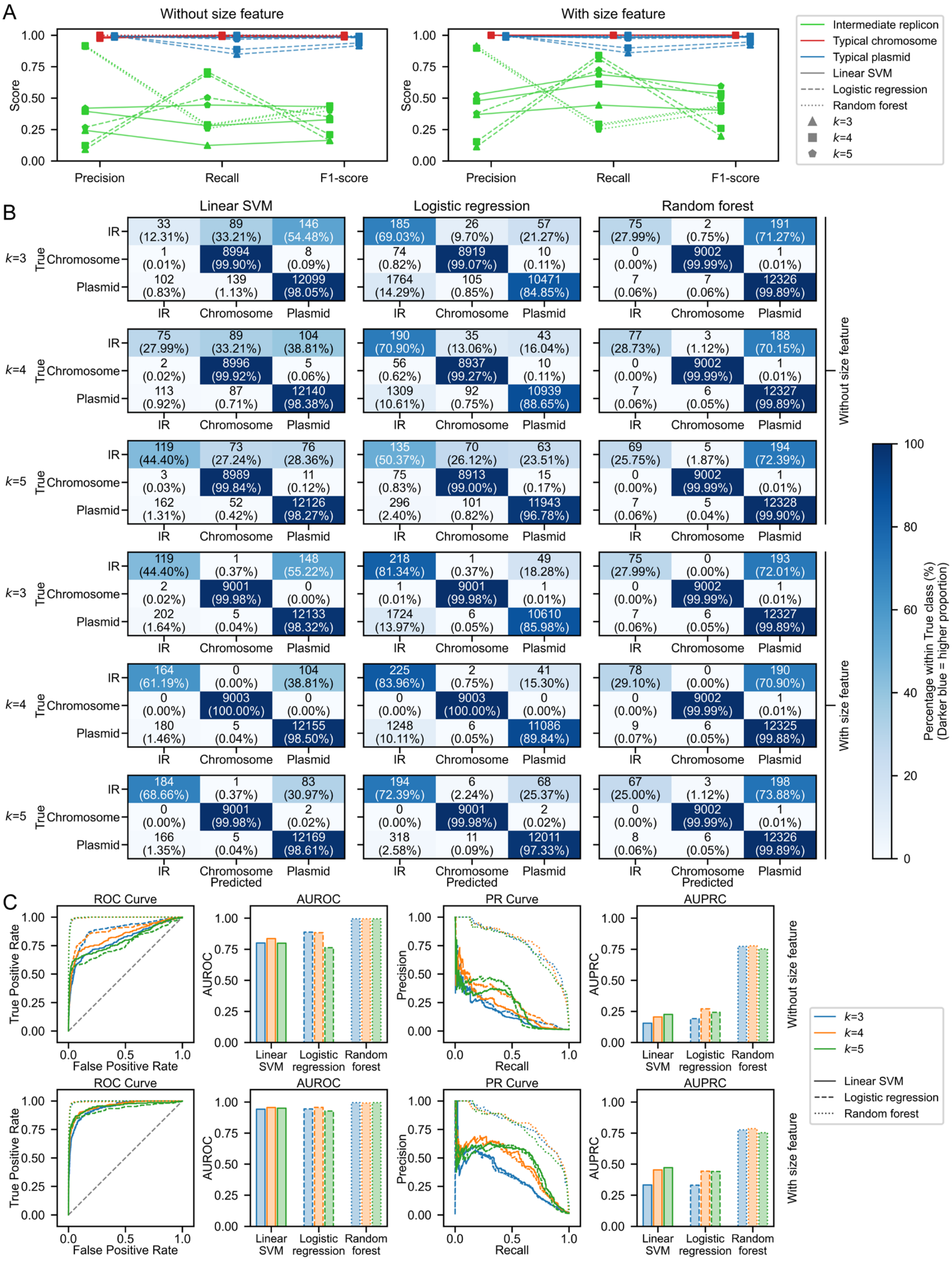


**Supplementary Fig. S17 | Machine-learning identification of intermediate replicons using integrated genomic features.**

(A) Precision, recall and F-score metrics of three classifiers (Linear SVM, Logistic regression, and Random forest). Classifiers were trained using replicon-level COG category frequencies and *k*-mer profiles ($k = 3$–5), with replicon size optionally included as an additional feature.

(B) Confusion-matrix results derived from classifier predictions. Percentage values represent the fraction of predicted entries within each true-label group.

(C) Receiver-operating-characteristic (ROC) curves, area-under-ROC (AUROC) bar plots, precision-recall (PR) curves, and area-under-PR-curve (AUPRC) bar plots for classifier-performance evaluation. Among the tested models, Random forest achieved the highest discriminative performance, and inclusion of replicon size further improved classification accuracy.

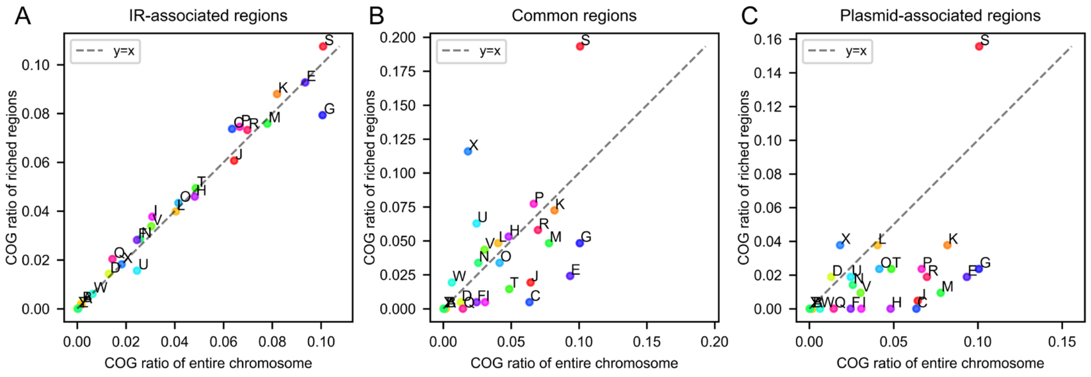


**Supplementary Fig. S18 | Comparison of COG proportions between distinct enriched regions and the entire chromosome.**

(A) COG-category ratios of IR-associated regions versus entire chromosomes.

(B) COG-category ratios of common regions versus entire chromosomes.

(C) COG-category ratios of plasmid-associated regions versus entire chromosomes.

The distribution of gene categories in regions associated with IRs was largely comparable to that of the whole chromosome. In contrast, the other two regions showed distinct functional profiles relative to the genomic background. For each panel, each dot corresponds to one COG category. The *x*- and *y*-values represent COG ratios calculated from whole chromosomes and the corresponding enriched regions, respectively. The dashed line indicates the $y=x$ reference line.

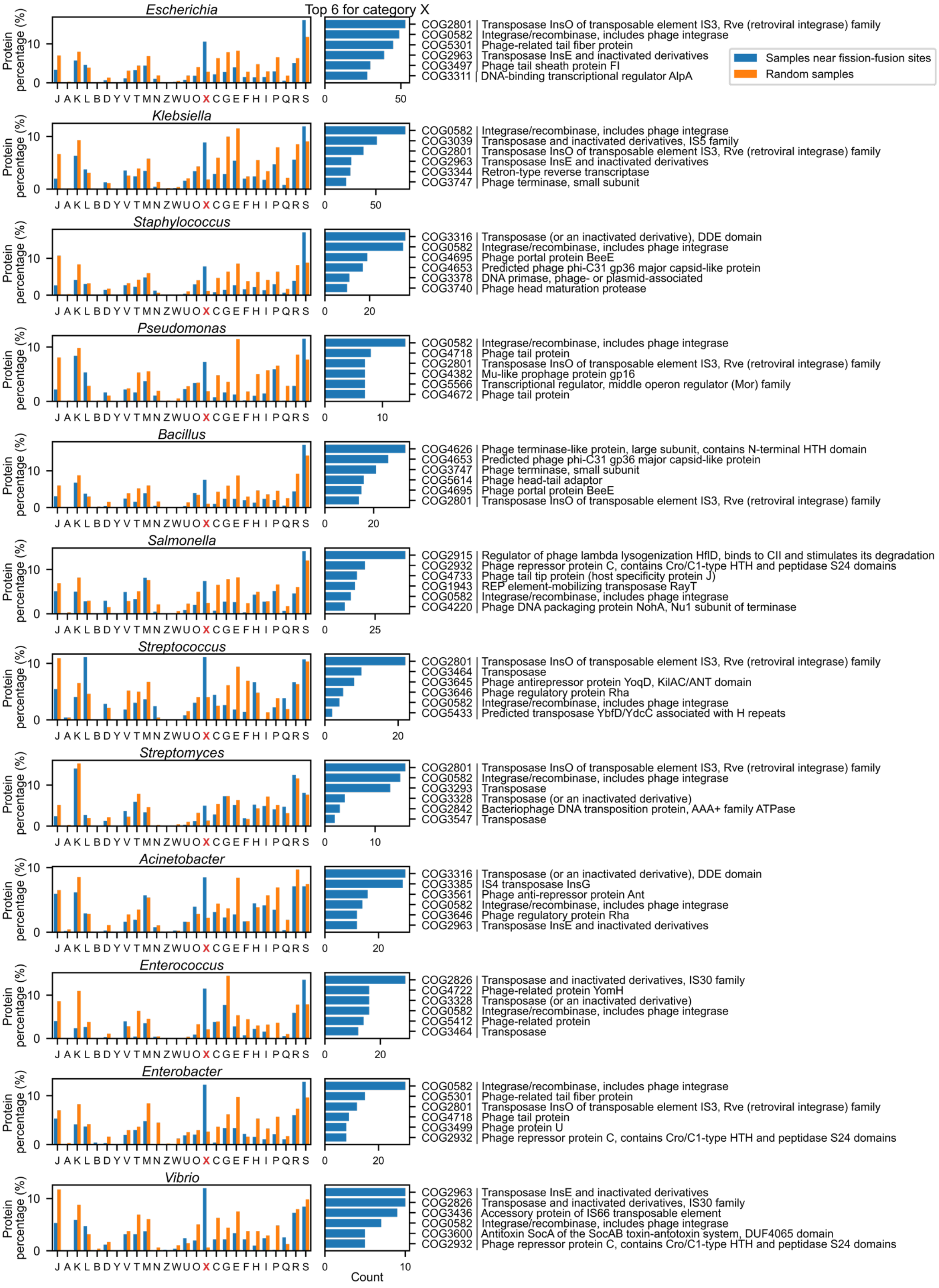


**Supplementary Fig. S19 | Functional features of genomic regions surrounding putative IR fission–fusion sites.** Protein functional categories enriched near putative IR fission–fusion sites were compared with length-matched randomly sampled chromosomal regions. Putative sites were defined based on the boundaries of chromosomal segments showing sequence homology to IRs. Flanking regions of 2.5 kb upstream and downstream were analyzed; only genera with more than 400 recovered CDSs are shown. Panels labelled “Top 6 for Group X” show the six highest-ranked

COG identifiers retrieved from samples near fission-fusion sites within category X; horizontal-axis values correspond to their respective count values.

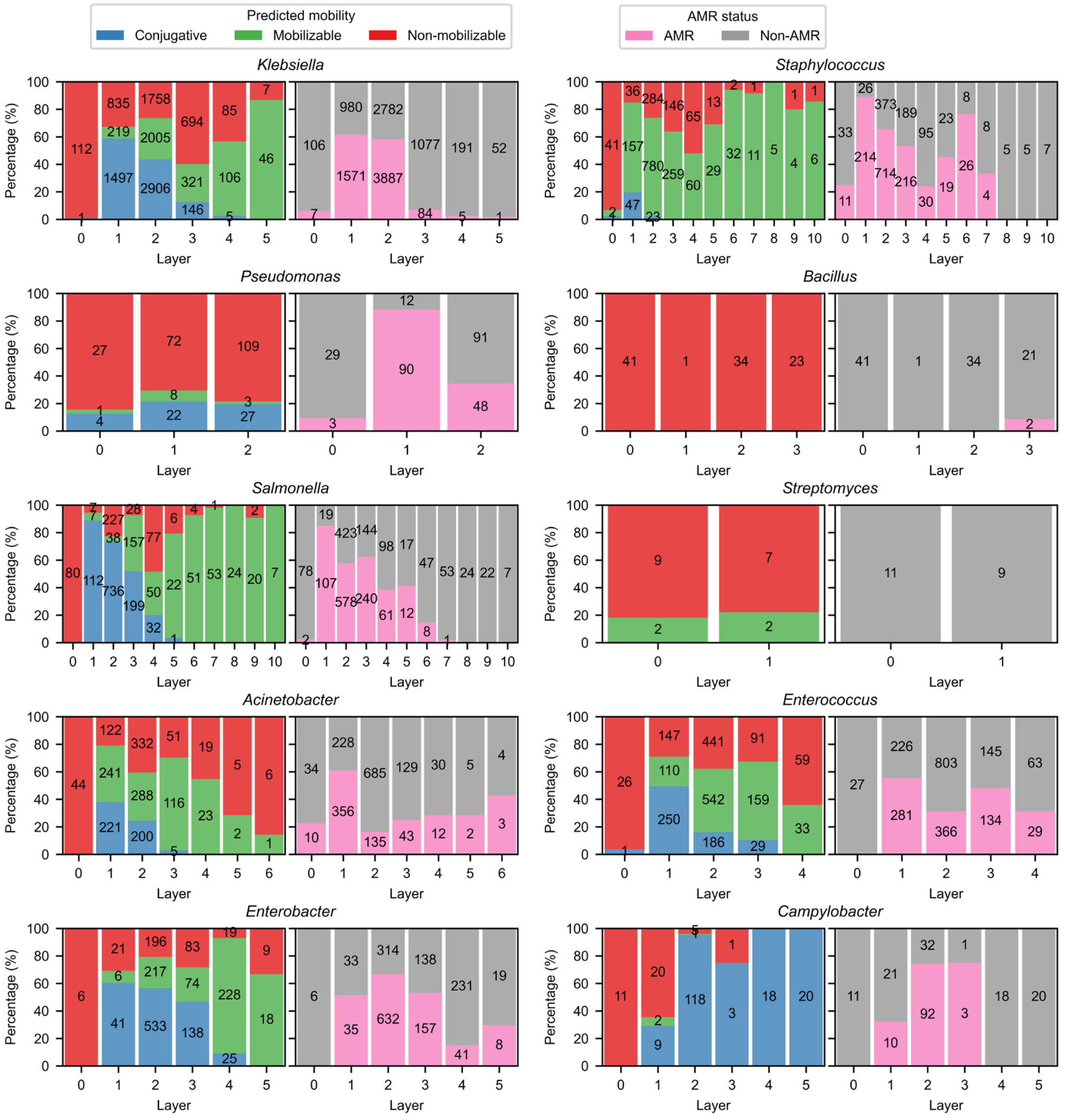


**Supplementary Fig. S20 | Enrichment of mobile and AMR-associated plasmids near intermediate replicons across bacterial genera.** Sequence association networks were constructed for genera containing both intermediate replicons (IRs) and typical plasmids. An edge between two replicons was defined if aligned regions covered at least the two-thirds of either replicon. IRs were assigned as layer 0, with typical plasmids organized by increasing network distance. Across genera, plasmids in proximal layers exhibited higher frequencies of conjugative traits and AMR associations.

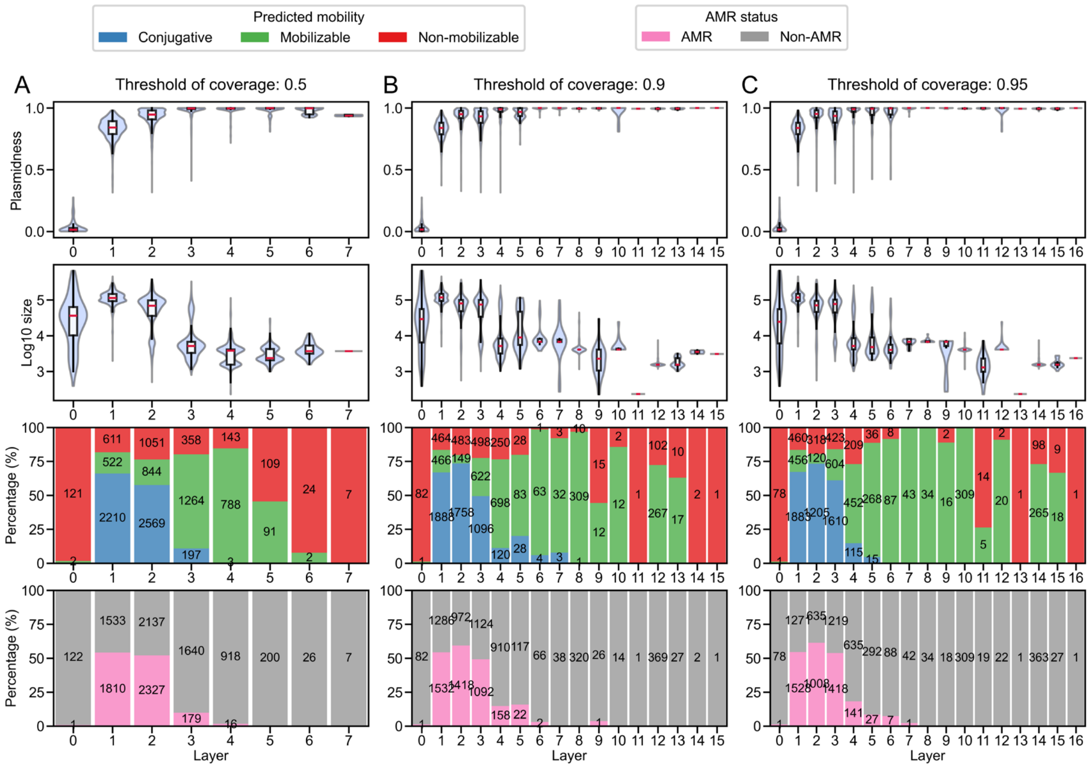


**Supplementary Fig. S21 | Statistical results of stratification under different coverage thresholds (*Escherichia*).**

(A) Distributions of plasmidness, $log_{10}$-transformed replicon size, predicted mobility and AMR status at a coverage threshold of 0.5.

(B) Distributions of plasmidness, $log_{10}$-transformed replicon size, predicted mobility and AMR status at a coverage threshold of 0.9.

(C) Distributions of plasmidness, $log_{10}$-transformed replicon size, predicted mobility and AMR status at a coverage threshold of 0.95.

Despite the use of different coverage thresholds, the four traits (plasmidness, size, mobility and AMR status) maintain consistent patterns with increasing layer number. An edge between two replicons was defined if aligned regions covered at least the coverage-threshold fraction of either replicon. Replicons were assigned into hierarchical network layers based on their shortest sequence-similarity distance to IRs (layer 0).

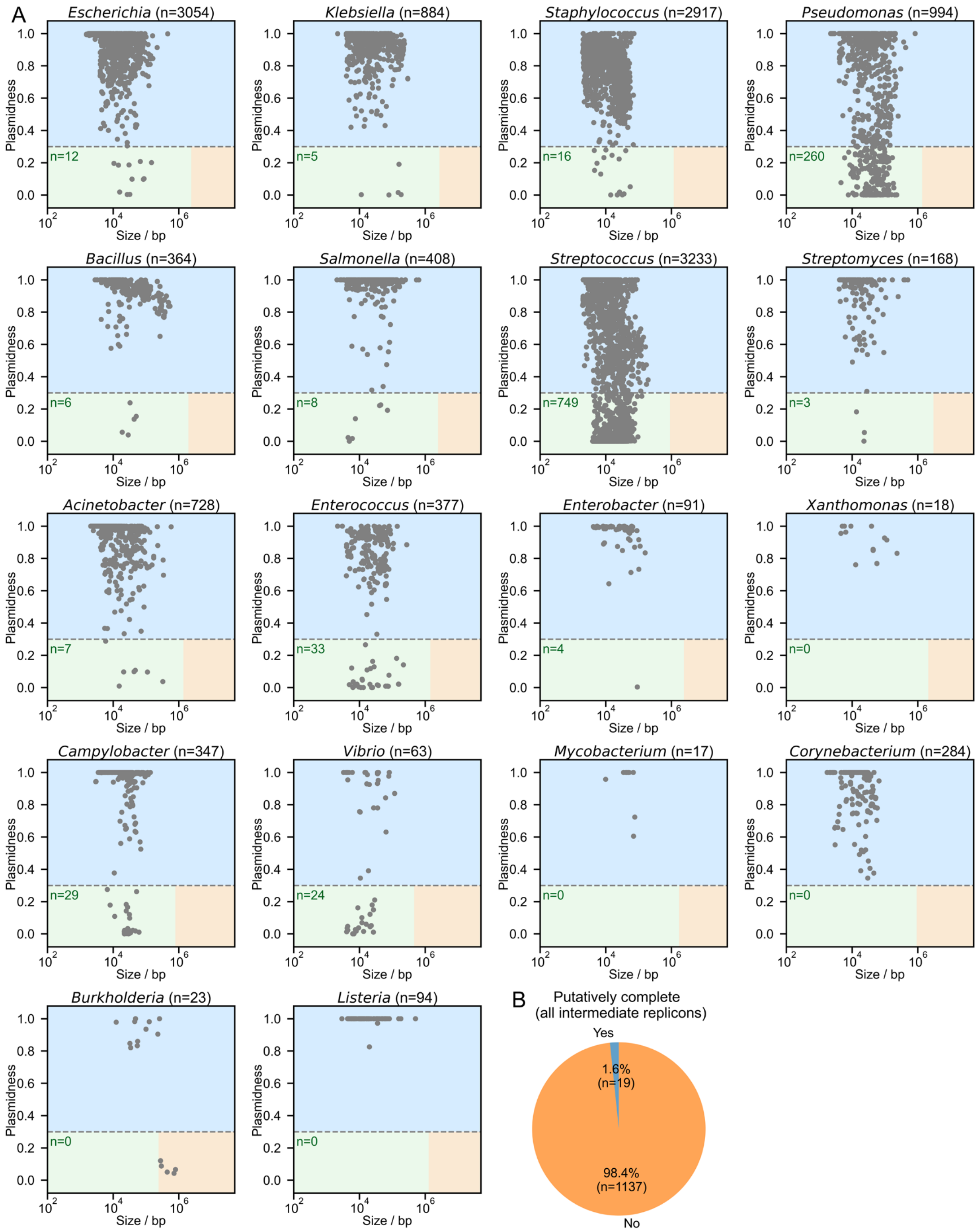


**Supplementary Fig. S22 | Detection of intermediate replicons among metagenome-assembled plasmid sequences from IMG/PR.**

(A) Scatter plots of plasmidness versus replicon size for representative bacterial genera. Plasmidness distributions were calculated for metagenome-assembled plasmid sequences across bacterial genera. In each panel, $n$ denotes the number of analyzed plasmid sequences, and the green label indicates the number of identified intermediate replicons.

(B) Pie chart illustrating the proportion of putatively complete intermediate replicons among all recovered intermediate replicons.
